# Myofibre-specific knockout of TGF-β type I receptors triggers muscle hypertrophy and promotes contraction and oxidative metabolism

**DOI:** 10.1101/2023.08.30.555466

**Authors:** A. Shi, M.M.G. Hillege, W. Noort, C. Offringa, G. Wu, T. Forouzanfar, W.M.H. Hoogaars, R.C.I. Wüst, R.T. Jaspers

## Abstract

Transforming growth factor-β (TGF-β) signaling is associated with progressive skeletal muscle wasting and fibrosis, while double knockout of TGF-β type I receptors *Acvr1b* and *Tgfbr1* results in hypertrophy. Gaining insights in how myofibre-specific knockout of these receptors affects muscle transcriptome, strength and mitochondrial activity could aid in the development of therapeutic interventions to improve muscle function. Here, we show that 3 months of myofibre-specific knockout of both receptors (dKO) in mice induced a 1.6-fold increase in gastrocnemius medialis mass and a 1.3-fold increase in maximal force. Soleus muscle mass and maximal force both increased 1.2-fold in dKO mice. Muscle hypertrophy in dKO mice was accompanied by a proportional increase in succinate dehydrogenase enzyme activity. Single receptor knockout caused minor phenotypical alterations. Transcriptome analyses revealed that gastrocnemius medialis had 1811 and soleus had 295 differentially expressed genes, mainly related to muscle contraction, hypertrophy, filament organization and oxidative metabolism. *Hgf* and *Sln* genes were strongly upregulated in both muscles of dKO mice, while *Sntb1* was downregulated. This in combination of transcriptional changes are associated with muscle hypertrophy and increased mitochondrial biosynthesis. Our study highlights that myofibre-specific interference with both TGF-β type I receptors concurrently stimulates myofibre hypertrophy and mitochondrial activity.

## INTRODUCTION

Muscle wasting diseases, including sarcopenia, cancer-induced cachexia, dystrophy, and physical inactivity, are associated with impaired physical performance which are largely due to reduced muscular strength and endurance ^1–9^. Optimal muscle force is determined by muscle physiological cross-sectional area (CSA) and the distribution of different types of myosin heavy chains, whereas muscle endurance is largely determined by the myofibre oxidative metabolic capacity, capillary density and myoglobin concentration ^10^. Typical features for muscle wasting disorders are the impairment of both muscle strength and fatigue resistance ^11,12^. Furthermore, in muscle wasting disorders, myofibres size is reduced, myofibres are lost and replaced by fibrotic tissue, and the force-generating capacity per CSA (i.e. specific force) and endurance capacity are decreased. It is therefore a clinical challenge to counterbalance these changes in muscle phenotype by concurrently stimulating myofibre hypertrophy and increasing myofibre oxidative capacity. Over a range of species, myofibre size has been shown to be inversely related to oxidative capacity ^13^, suggesting that both these traits are mutually exclusive. However, currently there is no effective treatment to restore muscle mass and aerobic metabolism simultaneously in muscle wasting disorders.

Interference with signalling pathways that affect muscle size and oxidative metabolism may ameliorate muscle dysfunction in many wasting conditions. Members of transforming growth factor-β (TGF-β) superfamily, such as TGF-β1, activin A, and myostatin, negatively regulate muscle mass and force, while they also contribute to muscle fibrosis ^14–17^. TGF-β1 also inhibits mitochondrial function and oxidative metabolism in both myoblasts and myotubes ^18^. A hallmark of myopathies and muscle wasting disorders is the elevated TGF-β signalling ^1,4,6^. Therefore, inhibition of TGF-β signalling could be a potential therapeutic approach to simultaneously increase muscle force and endurance.

TGF-β signalling is activated when type I receptors are phosphorylated and a complex is formed by its ligands and type II receptors. For the TGF-β type I receptor, TGF-β isoforms signal via TGF-β receptor type-1 (TGFBR1/ALK5), while activin A signals via activin receptor type-1B (ACVR1B/ALK4). Myostatin signals via both the TGFBR1 and ACVR1B ^19,20^. Phosphorylated type I receptors phosphorylate Smad2/3, resulting in the inhibition of muscle cell growth and differentiation ^21^.

Targeting one of the ligands or receptors of TGF-β1, myostatin and activin A increases protein synthesis and skeletal muscle mass. Intravenous administration of myostatin propeptide increased gastrocnemius medialis muscle mass by 20% 17 weeks after injection, while soleus muscle mass was not affected ^22^. Inhibition of type II receptor signalling of myostatin and activin A in mice by administration of neutralizing antibody for 5 weeks induced larger increases in muscle masses of gastrocnemius and plantaris (i.e. by 50%) than by using propeptide of myostatin (i.e. by 10%) ^23^. Moreover, targeting TGF-β type I receptors *Acvr1b* and *Tgfbr1* in mice has been shown to induce an increase in gastrocnemius muscle mass by about 200%, while targeting of type II receptors only increased muscle mass by only 50% ^24^. These findings indicate that targeting of TGF-β type I receptors stimulates muscle hypertrophy more pronouncedly than targeting of TGF-β family ligands or type II receptors. Noteworthy, the hypertrophy by interference with TGF-β signalling is much larger in type IIB myofibres than in type I myofibres which is presumably associated with higher expression levels of myostatin ^25^, type II receptors ^26^, and/or elevated *Pkm2* expression in fast-type myofibres ^27^. Although TGF-β signalling is well characterized, the mechanisms underlying the regulation of TGF-β1 receptors on muscle size, metabolisms and the muscle type-specific force-generating capacity are still unknown.

An increase in muscle mass does not necessarily imply a proportional increase in force-generating capacity. The myofibre hypertrophy by the knockout of myostatin in fast-type muscle is not proportionally associated with an increase in force-generating capacity, hence muscle specific force is reduced ^26,28,29^. Although it has been reported that the reduced specific force in fast-type muscle of myostatin-null mice was associated with an increased myonuclear domain size, reduced muscle stiffness ^30^ and diminished expression of collagen genes ^31^, the underlying molecular mechanism is unclear. Whether disproportional increase in force-generating capacity and myofibre size is attributable to the changes in transcript levels of muscle contractile components, and/or cell-matrix adhesion proteins requires a thorough investigation. Whole transcriptome analysis of both fast-and slow-type myofibres lacking TGF-β type I receptors may provide insight in the signalling pathways involved.

Another phenotypical characteristic of myofibres is the oxidative metabolic capacity which is a key determinant of the maximal sustainable muscle power ^32^. Several studies have shown that muscles of myostatin-null mice show increased cytochrome c oxidase enzyme activity and elevated *Pgc1a* gene expression levels ^33,34^. In line with this, TGF-β1 supplementation reduced mitochondrial complex IV protein abundance in both myoblasts and myotubes *in vitro* ^18^. These findings suggest that blockade of TGF-β signalling by targeting type I receptors inhibiting myostatin and TGF-β1 signalling may concurrently increase muscle size and mitochondrial activity, and as such could be an effective treatment in muscle wasting disorders.

The aim of this study was to investigate the effects of targeting TGF-β type I receptors on (1) fast-and slow-type muscle phenotypes, (2) the force-generating capacity and the underling molecular mechanisms, (3) muscle oxidative metabolism. For this purpose, type I receptors *Acvr1b* and *Tgfbr1* were knocked out individually and simultaneously (dKO). We assessed the transcriptome, *in situ* force-generating capacity *in situ*, and oxidative metabolism of both glycolytic, low oxidative gastrocnemius medialis and high oxidative soleus muscles. Here we show that targeting either *Acvr1b* or *Tgfbr1* individually did not significantly increase muscle mass and force. In contrast, the myofibre-specific lack of both type I receptors resulted in substantial alterations in expression levels of genes related to muscle growth, contraction, myoblast differentiation, filament organization, connective tissue remodelling, and metabolism. Both gastrocnemius and soleus muscles of dKO mice were larger and stronger, while specific force was reduced in gastrocnemius. Strikingly, simultaneous knockout of type I receptors showed concurrent sizable myofibre hypertrophy and increased oxidative metabolism.

## RESULTS

### Fast-and slow-type muscles lacking both TGF-β type I receptors show different transcriptional profiles

To genetically inactivate *Acvr1b* and *Tgfbr1* in a myofibre-specific manner, tamoxifen-induced Cre recombinase was used in six-week-old HSA-MCM:*Acvr1b^fl/fl^,* HSA-MCM:*Tgfbr1^fl/fl^,* HSA-MCM: *Acvr1b^fl/fl^:Tgfbr ^fl/fl^* mice. Cre positive HSA-MCM mice were injected with tamoxifen as the control group (**Figure 1. A**). Animals were sacrificed after three months (**Figure 1. B**). Gene expression levels of *Acvr1b* in gastrocnemius were significantly reduced in gastrocnemius of *Acvr1b* KO mice and dKO mice compared to those of control. Gene expression levels of *Tgfbr1* were reduced in gastrocnemius of the *Tgfbr1* KO mice and dKO mice compared to those of control mice. For soleus muscle, although RNA-seq and q-PCR did not show differences in gene expression levels of *Acvr1b* or *Tgfbr1* between groups (**Figure 1. C**, **D**), spatial quantification of expression levels of *Acvr1b* and *Tgfbr1* using ISH of RNA scope confirmed the reduced number of *Acvr1b/Tgfbr1* mRNA per nuclear in both gastrocnemius and soleus of *Acvr1b* KO/*Tgfbr1* KO mice and dKO mice (**Supplementary Figure 5**). Both muscle mass and maximal contractile force were increased in soleus muscles of dKO mice which indicates that muscle function and phenotype were altered by the knockout of both *Acvr1b* and *Tgfbr1*. Body mass of dKO animals was increased progressively and reached a plateau at ∼7 weeks after tamoxifen injection, while the other two groups showed typical growth curves that were similar to that of the wild type (**Figure 1. E**). Both gastrocnemius and soleus muscle mass of the dKO mice were higher than those of the other groups (**Figure 1. F**, **G** and **Supplementary Table 1**). RNA-seq analysis revealed that major alterations in gene expression were in gastrocnemius of dKO mice but not in muscles lacking either *Acvr1b* or *Tgfbr1* (**Figure 1. H**, **I**). For gastrocnemius of dKO mice 1811 genes were differentially expressed (enriched by Deseq2 with a false discovery rate *P* < 0.05 and fold change ≥ 1.5) whereas dKO soleus muscles showed 295 differentially expressed genes (DEGs). Many DEGs in dKO muscles were not observed in muscles of single receptor knockout (**Figure 1. J**, **Supplementary Figure 1. A, B and Supplementary Table 2**, **3**). Gene ontology (GO) enrichment analysis revealed that pathways related to muscle contraction, growth, filament organization, stem cell differentiation, matrix remodelling, and metabolism were predominantly altered in gastrocnemius muscles of dKO mice, rather than in soleus muscle (**Figure 1. K**).

**Figure 1.**
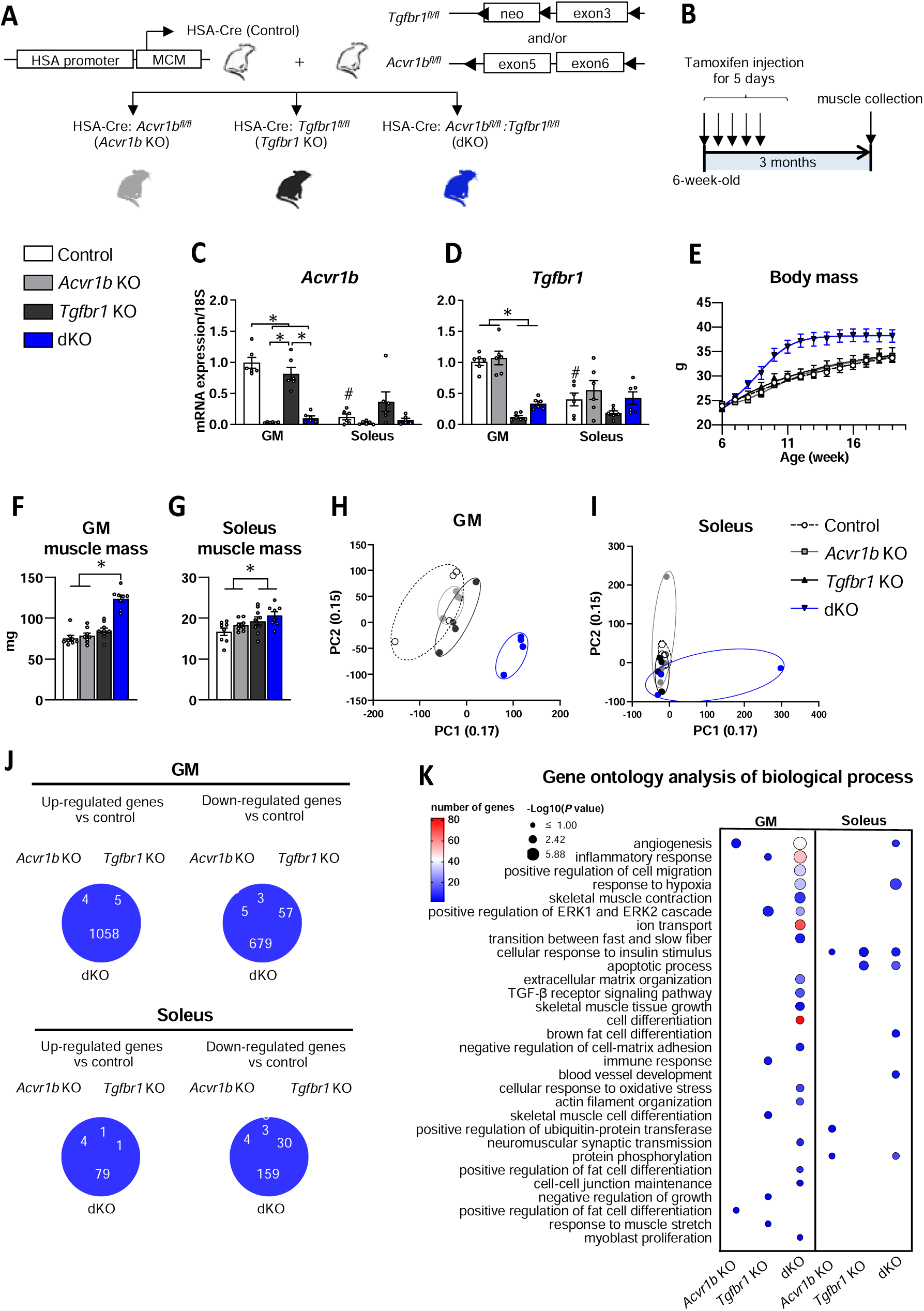
Simultaneous skeletal muscle-specific TGF-β type I receptors knockout induces muscle hypertrophy and transcriptomic change. (**A**) Experiment design of skeletal muscle-specific knockout of TGF-β type I receptors in mice. (**B**) Schematic of the study outline indicates the time points for gene knockout and muscle collection. Gene expression of (**C**) *Acvr1b* and (**D**) *Tgfbr1* (n=6). (**E**) Body mass from onset of knockout until sacrifice (n=7-8). Muscle mass of (**F**) gastrocnemius (GM) and (**G**) soleus. Principal component analysis (PCA) of (**H**) gastrocnemius and (**I**) soleus muscles (n= 4). (**J**) Venn diagram of up and down-regulated differentially expressed genes (DEGs) in gastrocnemius and soleus of knockout mice (enriched by DESeq2 with a false discovery *P* < 0.05 and fold change ≥ 1.5). (**K**) Bubble plot of gene ontology enrichment analysis on biological processes in gastrocnemius and soleus between control and gene knockout mice (false discovery *P* < 0.05 and fold change ≥ 1.5). Data are shown as mean ±DSEM. *: *P* < 0.05. #: *P* < 0.05 compared to GM of control mice. Two-way ANOVA with Bonferroni post hoc test was used to analyse data. Independent t-tests were used to compare data for GM and soleus of control group. See also **Supplementary Figure 1**.

### Knockout of TGF-β type I receptors in gastrocnemius and soleus muscles differentially affects muscle force-generating capacity

RNA-seq analysis showed that TGF-β type I receptor knockout in both gastrocnemius and soleus muscles induced DEGs which were associated with skeletal muscle contraction (**Figure 2. A**). For dKO gastrocnemius, expression levels of genes that encode contractile filaments were decreased. In particular, dKO gastrocnemius showed lower expression levels of genes encoding thick filament-associated proteins (*Mylk2, Myl2, Myh7*) and slow-myofibre type troponin and tropomyosin (*Tnnc1, Tnni1, Tnnt1, Tpm3*). Besides, gastrocnemius of dKO mice had decreased gene expression levels of genes involved in protein degradation (*Foxo1*, *Foxo4*) (**Supplementary Data**). In addition, targeting both type I receptors in gastrocnemius increased gene expression levels of giant sarcomeric protein (*Obsl1*) and proteins modulating contractility such as *Mybph* and *Actc1*. Genes encoding acetylcholine receptors subunit proteins of the postsynaptic neuromuscular junction (*Chrna1*, *Chrng*, *Chrnd*) were more expressed in gastrocnemius than in soleus muscle of dKO mice (**Figure 2. A**, **Supplementary Data**). Based on these altered gene expression landscapes, we next studied the muscle functional contractile properties. **Figure 2. B**-**E** show typical examples of twitch and tetanus force tracings of a gastrocnemius and a soleus of a dKO and a control mouse. The lack of *Acvr1b* significantly increased maximal isometric tetanic force in gastrocnemius. Post hoc analysis showed that the maximal tetanic force of gastrocnemius in dKO mice (2123.46±142.71 mN) was 1.3-fold higher than that of control mice (1669.65±105.47 mN) (**Figure 2. F**). The specific force of gastrocnemius in dKO mice was 77% (17.25±1.19 mN/mg) of that of control mice (22.32±1.45 mN/mg) (**Figure 2. H**), indicating disproportional increases in muscle size and force. Maximal tetanic force in soleus muscle of dKO mice was higher compared to that in *Acvr1b* KO and control animals (**Figure 2. G**), while specific force was unaltered (**Figure 2. I**), indicating that the increase in force-generating capacity was proportional to the increase in mass. Passive force of both muscles at optimal length did not differ between groups (**Figure 2. J**, **K**). Force-frequency relations were assessed to determine whether muscle calcium sensitivity was affected by TGF-β receptor knockout. A slight leftward shift in force-frequency relation was observed in gastrocnemius lacking *Tgfbr1*, however it did not occur in soleus (**Figure 2. L**, **M**).

**Figure 2.**
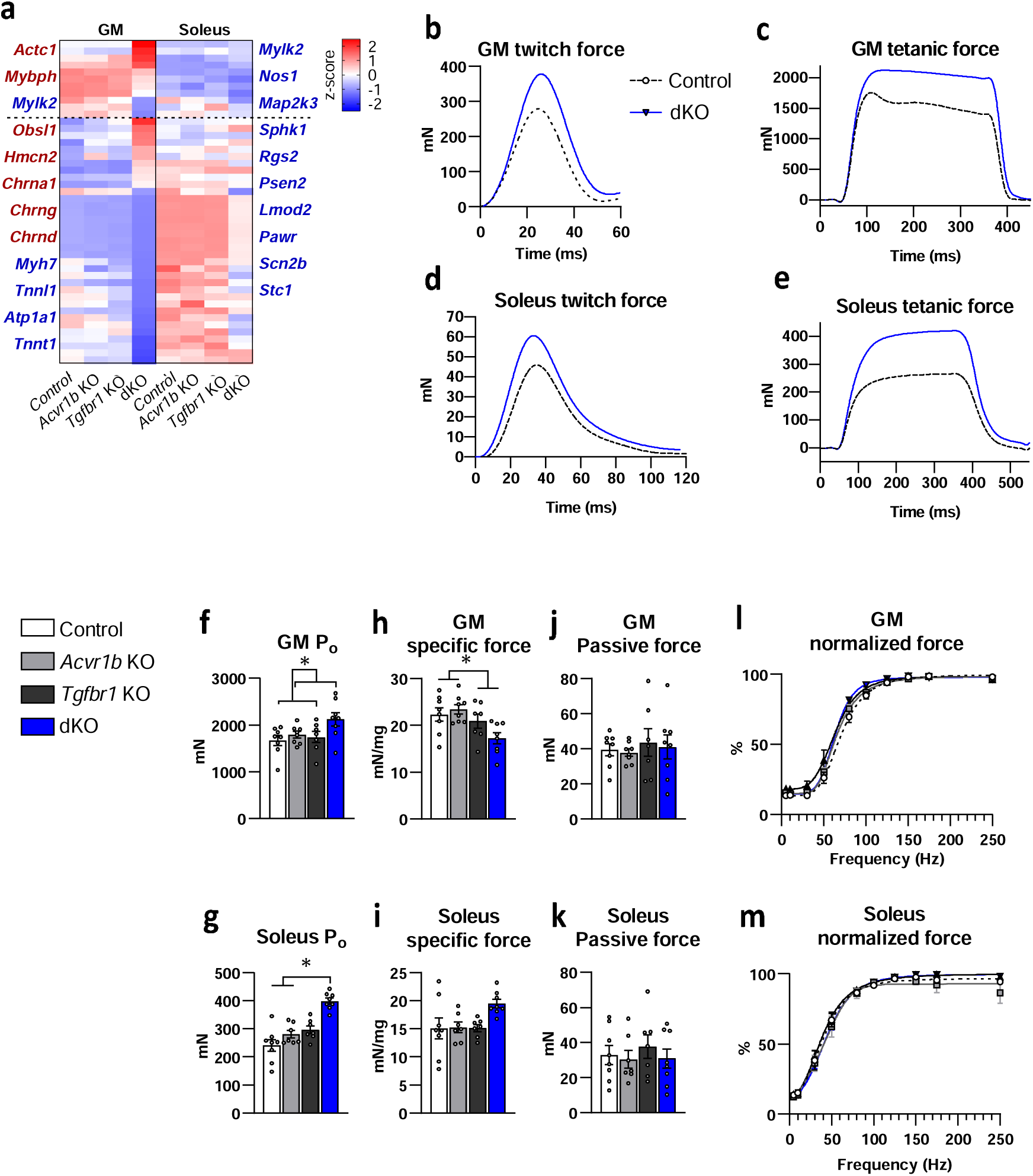
The lack of TGF-β type I receptors affects isometric force generation. (**A**) Heatmap of genes in the cluster of skeletal muscle contraction that exemplifies DEGs in gastrocnemius and soleus muscles of dKO mice were depicted on the left and right side of heatmap, respectively (n = 4). Upregulated genes were in red, while downregulated genes were in blue. Typical examples of twitch and tetanus force tracings of a (**B**), (**C**) gastrocnemius and (**D**), (**E**) soleus muscle of a dKO and a control mouse. Maximal isometric tetanic force (P_o_) of (**F**) gastrocnemius and (**G**) soleus muscles (n= 7-8). Specific force of (**H**) gastrocnemius and (**I**) soleus muscles (n = 7-8). Passive force of P_o_ in (**J**) gastrocnemius and (**K**) soleus muscles (n=7-8). The force-frequency relations of (**L**) gastrocnemius and (**M**) soleus muscles (n=7-8). Data are shown as mean ±DSEM. *: *P* < 0.05. Two-way ANOVA with Bonferroni post hoc tests were performed to analyse data. Also see **Supplementary Table 4**.

To determine effects on excitation-contraction coupling, contractile twitch characteristics of gastrocnemius and soleus muscles were determined. Gastrocnemius twitch characteristics did not differ between groups (**Supplementary Table 4**). The maximal twitch force of dKO soleus was significantly higher compared to that of control mice. No significant differences were observed in twitch/tetanus ratio, time-to-peak twitch tension, normalised maximum rise in tension or half-relaxation time between groups. Together, these results indicate that force-generating capacity of muscle lacking both *Acvr1b* and *Tgfbr1* was increased, albeit fast-type muscle showed a disproportional increase in muscle mass and force-generating capacity.

### Simultaneous knockout of TGF-β type I receptors increases myofibre size and induces transcriptional changes related to sarcomere organization and transmembrane complex

Since muscle force was increased after knockout of both TGF-β type I receptors, we assessed muscle histological, cellular and molecular determinants of muscle phenotype. In the high oxidative region of gastrocnemius muscle, myofibre type distribution of dKO mice was shifted from type I and IIA myofibres to IIX and IIX/IIB compared to that in *Acvr1b* KO animals (**Figure 3. A**-**C**, **Supplementary Figure 2**), while myofibre type distribution was not different in muscles of single knockout mice. FCSA of dKO animals was 1.5-fold (3584±193 μm^2^) and 3.2-fold (10651±803 μm^2^) larger in the high and low oxidative regions of gastrocnemius muscle, respectively, compared to that of the other groups (**Figure 3. E**-**F**). The number of myonuclei per myofibre in high oxidative region of gastrocnemius in dKO mice was increased with the increase in FCSA, while in the low oxidative region of gastrocnemius in dKO mice, nuclear number per myofibre was similar as that in control, resulting in 2.9-fold larger myonuclear domain compared to that in control muscles (**Figure 3. H**, **I**). For soleus muscles of dKO mice, myofibre type I/IIA percentage was slightly increased in soleus (**Figure 3. D**) and FCSA (2822±216 μm^2^) was ∼1.5-fold higher (**Figure 3. G**). Myonuclear number and myonuclear domain in soleus muscle of dKO mice were not different from those in control (**Figure 3. H**, **I**).

**Figure 3.**
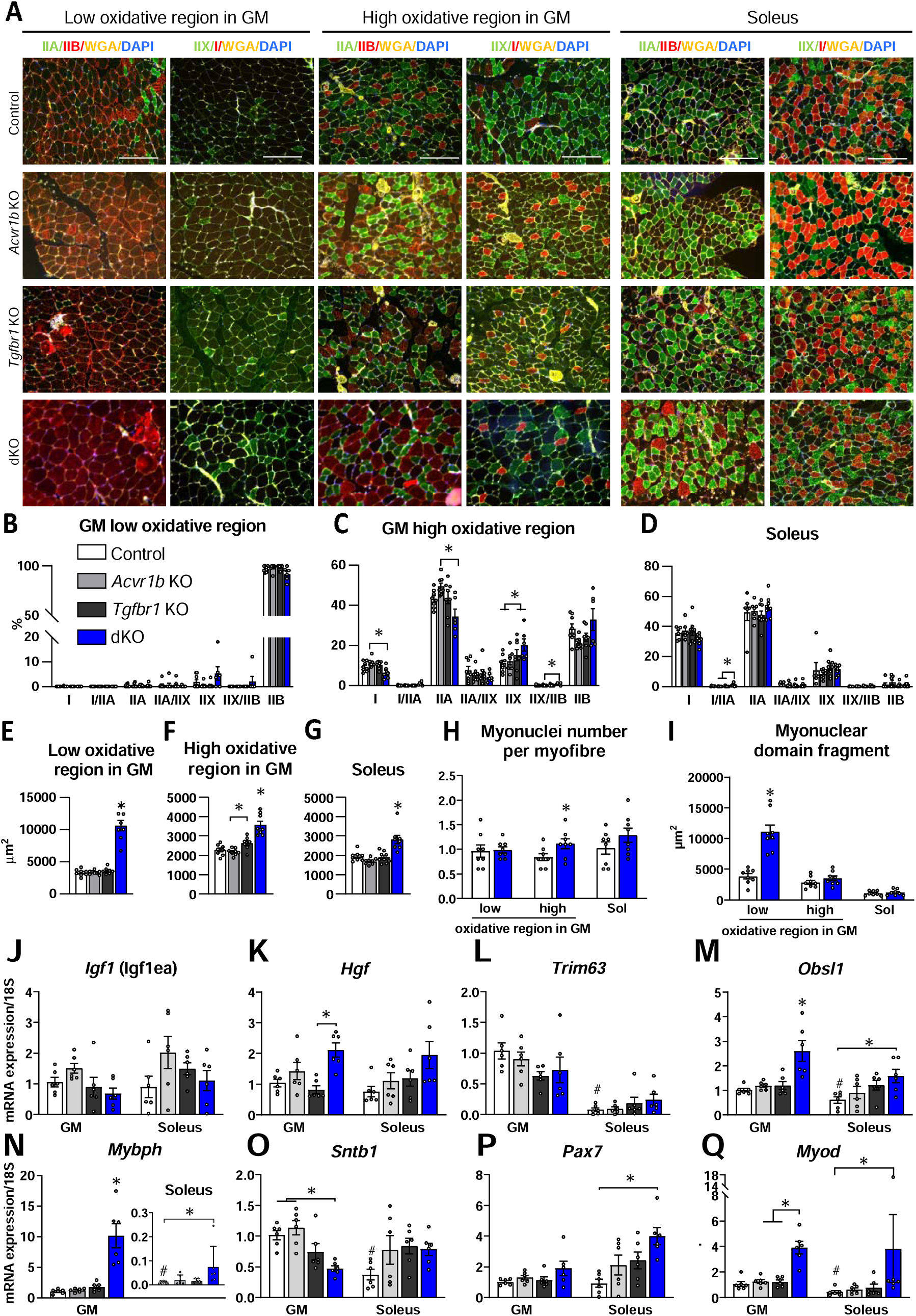
Knockout of TGF-β type I receptors affects muscle morphology and gene expression. (**A**) Immunofluorescence (IF) staining of type IIA (green), IIB (red), I (red), and IIX (green) myosin heavy chain in gastrocnemius and soleus muscles. Yellow: wheat germ agglutinin (WGA), blue: DAPI. Scale bar=250μm. Myofibre type distribution in (**B**) low, (**C**) high oxidative region in gastrocnemius, and (**D**) soleus. Myofibre cross-sectional area in (**E**) low, (**F**) high oxidative region in gastrocnemius, and (**G**) soleus muscles (n=7-8). (**H**) Myonuclei number per myofibre and (**I**) myonuclear domain in control and dKO mice (n=7-8). qPCR for (**J**) *Igf1ea*, (**K**) *Hgf,* (**L**) *Trim63,* (**M**) *Obsl1*, (**N**) *Mybph*, (**O**) *Sntb1*, (**P**) *Pax7* and (**Q**) *Myod* (n=6). Two-way ANOVA with Bonferroni post hoc tests were performed to analyse data. Independent t-tests were used to compare GM and soleus data of the control group. Data are shown as mean ±DSEM. *: *P* < 0.05. A single star indicates a significant difference compared to all other groups of the same muscle. #: *P* < 0.05 compared to GM of control mice. Also see **Supplementary Figure 2**.

We hypothesized that the muscle hypertrophy in dKO mice was caused by higher rates of protein anabolism and reduced catabolism. Hepatocyte growth factor (*Hgf)* expression was increased in gastrocnemius muscles in dKO mice compared to that in *Tgfbr1* KO mice (**Figure 3. K**). However, expressions levels of genes encoding protein synthesis and protein degradation were not affected, including insulin-like growth factor 1 (*Igf1ea*) (**Figure 3. J**), mechano growth factor (*Igf1*ec) (**Supplementary Figure 1. D**) and E3 ubiquitin protein ligase (*Trim63*) (**Figure 3. L**).

Next, using qPCR we investigated whether the altered force-generating capacity in dKO mice was associated with the change in expression level of genes encoding proteins in organization of contractile structures. Gene expression levels of *Obsl1* and *Mybph* were increased in both gastrocnemius and soleus muscle of dKO mice (**Figure 3. M**, **N**), while those of *Ttn*, *Obscn* and *Ankrd2* were not different between groups (**Supplementary Figure 1. D**-**F**), suggesting adaptation of sarcomere architecture in hypertrophic muscle. Gene expression levels of scaffold proteins that interact with dystrophin glycoprotein complex (DGC) were assessed. DGC plays a major role in maintaining membrane integrity and interaction of sarcolemma to extracellular matrix (ECM) ^35^. Although gene expression level of δ-sarcoglycan (*Sgcd*) was not changed (**Supplementary Figure 1. G**), β1-syntrophin (*Sntb1*) expression levels were substantially lower in dKO gastrocnemius compared to those in control mice but not in soleus muscles (**Figure 3. O**). Reduced levels of *Sntb1* expression may interrupt force transmission from the sarcomeres to the ECM during muscle contraction. In addition, gene expression levels of adenosine A1 receptor (*Adora1*) and calcium voltage-gated channel subunit alpha1 S (*Cacna1s*) did not differ between control mice and both muscles lacking either *Acvr1b* or *Tgfbr1*, indicating limited effects of blocking type I receptors on presynaptic neuromuscular junction and sarcoplasmic reticulum calcium release (**Supplementary Figure 1. H**, **I**). In addition, sarcolipin expression (*Sln*) showed a 20-fold increase in gastrocnemius and soleus muscles of dKO mice (**Supplementary Data**). Moreover, expression of stem cell-specific transcription factor *Pax7* was increased in soleus and the early marker for myogenic commitment, *Myod*, was increased in both gastrocnemius and soleus muscles of dKO mice (**Figure 3. P**, **Q**). Although muscle mass was increased substantially in dKO mice, RNA concentration (ng/mg) was not different compared to other groups (**Supplementary Figure 1. J**). Together, these data indicate that gastrocnemius muscles without both *Acvr1b* and *Tgfbr1* show a substantial larger myofibre size without accretion of nuclei and show a shift towards a more fast-twitch myofibre phenotype. The myofibre hypertrophy was associated with increased gene expression of *Hgf* and altered gene expression related to sarcomeric organization and transmembrane complexes.

### Eccentric force is reduced during serial eccentric contractions in gastrocnemius lacking both TGF-β type I receptors despite an increase in connective tissue content

Since the lack of dystrophin of DGC in muscle has been shown to enhance muscle susceptibility to injury in muscle ^36^, it remains unclear whether reduced expression of another component of the DGC, the *Sntb1,* in dKO mice caused a decline in tetanic force during a series of eccentric contraction. We show that in the absence of both type I receptors, eccentric force-generating capacity was better preserved in soleus than in gastrocnemius. For gastrocnemius of dKO animals, the eccentric force curve was significantly lower than for muscles of the other groups (**Figure 4. A**). For soleus, eccentric force curves were not different between groups (**Figure 4. B**). After 15 eccentric contractions, in both muscles, the decline in maximal isometric tetanic force compared to the initial P_o_ prior to the series of eccentric contractions did not differ between groups (**Supplementary Figure 3**). Since muscle connective tissue plays a role in protecting muscle against injury ^37,38^, we then tested whether the decline in force during eccentric contractions was associated with a relatively lower content of connective tissue in the muscles of dKO mice.

**Figure 4.**
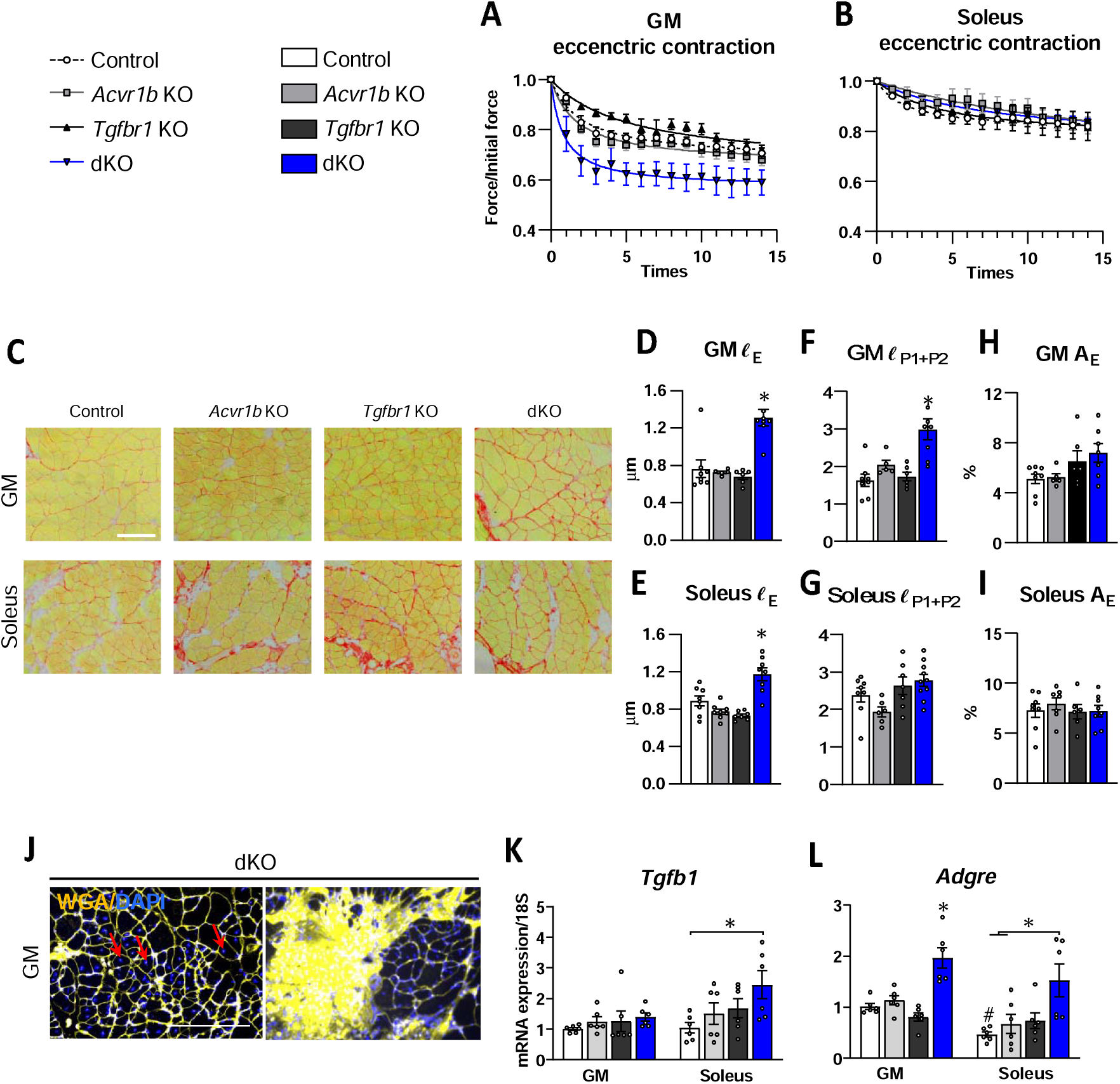
Muscle-specific TGF-β type I receptors deficiency affects force of during serial eccentric contraction and connective tissue content. Fold change of muscle force relative to initial contraction in 15 consecutive eccentric contractions in (**A**) gastrocnemius (GM) and (**B**) soleus (n=7-8). (**C**) Sirius red staining of gastrocnemius and soleus muscle. Scale bar=100μm. (**D**)-(**E**) Endomysium thickness (ℓ_E_) of gastrocnemius and soleus muscles. (**F**)-(**G**) Primary and secondary perimysium thickness (ℓ_P1+P2_) of gastrocnemius and soleus. (**H**)-(**I**) Endomysium area percentage per myofibre cross-sectional area (A_E_) of gastrocnemius and soleus (n=5-8). (**J**) Myofibres with central nuclei (red arrows) and excessive connective tissue deposition in the low oxidative region of gastrocnemius in dKO animals. Scale bar=250μm. qPCR expression levels for (**K**) *Tgfb1* and (**L**) *Adgre* (n=6). Two-way ANOVA with Bonferroni post hoc tests were performed to analyse data. Independent t-tests were used to compare qPCR data of gastrocnemius and soleus muscle of control groups. Data are shown as mean ±DSEM. *: *P* < 0.05. A single star indicates a significant difference compared to other groups of the same muscle. #: *P* < 0.05 compared to GM of control mice.

In both gastrocnemius and soleus muscles of dKO mice, endomysium was thicker than in those muscles of control mice (**Figure 4. C**-**E**). Thickness of primary and secondary perimysium in dKO gastrocnemius was increased compared to control, but not in soleus muscle (**Figure 4. F**, **G**). However, the percentages of endomysium area per FCSA in both gastrocnemius and soleus muscles of each genotype were not different from those in control (**Figure 4. H**, **I**). Note that excessive connective tissue deposition and small regions of myofibres with central nuclei were found in most low oxidative regions of gastrocnemius of dKO mice, but not in soleus (**Figure 4. J**). These regions are indicative of higher regenerative activities of cells in dKO fast-type muscle. We expreted that the increased ECM deposition and the regenerating myofibres in gastrocnemius of dKO mice were accompanied by an increase in gene expression related to the pro-fibrotic and inflammation. In contrast, in the absence of both *Acvr1b* and *Tgfbr1*, *Tgfb1* expression was not increased in gastrocnemius but was increased in soleus muscles (**Figure 4. K**). Gene expression of *Adgre,* which encodes the macrophage marker protein F4/80, was upregulated in both gastrocnemius and soleus muscles of dKO mice (**Figure 4. L**). These results indicate that simultaneous receptors knockout of both TGF-β type I receptors in both gastrocnemius and soleus muscles promotes ECM deposition proportionally with the increase in myofibre size. The decline in eccentric contractile force of fast-type gastrocnemius of dKO mice may be associated with the presence of newly formed myofibres in these muscles rather than with the change in ECM.

### Simultaneous receptors knockout increases integrated succinate dehydrogenase (SDH) activity and maintains mitochondrial enzyme activity in both gastrocnemius and soleus muscles

Muscle oxidative phosphorylation capacity relates inversely to myofibre size over a range of specie. A high mitochondrial enzyme activity and a larger myofibre size are mutually exclusive ^13,39^. To study whether TGF-β signalling affects mitochondrial oxidative metabolism, we hierarchically clustered genes in the transcriptome that were associated with oxidative metabolism from RNA-seq analysis. Expression levels of genes related to intracellular oxygen transport and mitochondrial enzyme activity were reduced in both muscles of dKO mice by RNA-seq, i.e. myglobin (*Mb*) and peroxisome proliferator-activated receptor gamma coactivator 1-alpha (*Ppargc1a*) (**Figure 5. A**, **Supplementary Data**). SDH activity was quantified to obtain an estimate of the oxidative capacity (**Figure 5. B**). The integrative SDH activity is the SDH activity times FCSA, which indicates total oxidative enzyme activity per myofibre per FCSA. For gastrocnemius, SDH activities in both low and high oxidative regions in *Tgfbr1* KO animals were higher than those in *Acvr1b* KO mice (**Figure 5. C**, **D**). In the low oxidative region of gastrocnemius, integrated SDH activity of dKO mice was higher than that in the other three groups (**Figure 5. C**). In the high oxidative regions of gastrocnemius, integrated SDH activity was higher in dKO mice than that in control and *Acvr1b* KO mice, but did not differ from *Tgfbr1*KO mice (**Figure 5. D**). For soleus, SDH activity in *Tgfbr1* KO mice was higher compared to that in all other groups (**Figure 5. E**). Integrated SDH activity in *Tgfbr1* KO and dKO mice was increased compared to that in *Acvr1b* KO mice (**Figure 5. E**). We observed a substantial rightward shift in the relationship between FCSA and SDH in *Tgfbr1* KO and dKO mice (**Figure 5. F**). This indicates that knockout of *Tgfbr1* increases muscle oxidative capacity and that SDH activity within myofibres is higher and independent of the hypertrophy in dKO mice. q-PCR was performed to further determine the expression levels of genes encoding mitochondrial respiratory chain enzymes and proteins involved in angiogenesis. Despite reduced gene expression shown by RNA-seq, q-PCR could not show changes in expression levels of *Sdhb*, *Mtco2*, *Ppargc1a* and *Vegfa* in either gastrocnemius or soleus of dKO mice compared to control mice (**Figure 5. G**-**J**). In addition, cytochrome c oxidase activity per volume cytoplasm in of both muscles in dKO mice was not different from control mice (**Figure 5. K**, **Supplementary Figure 4. A**). Together with the unaltered SDH activity, these findings indicate that mitochondrial biosynthesis capacity was enhanced in each hypertrophic myofibre of dKO mice.

**Figure 5.**
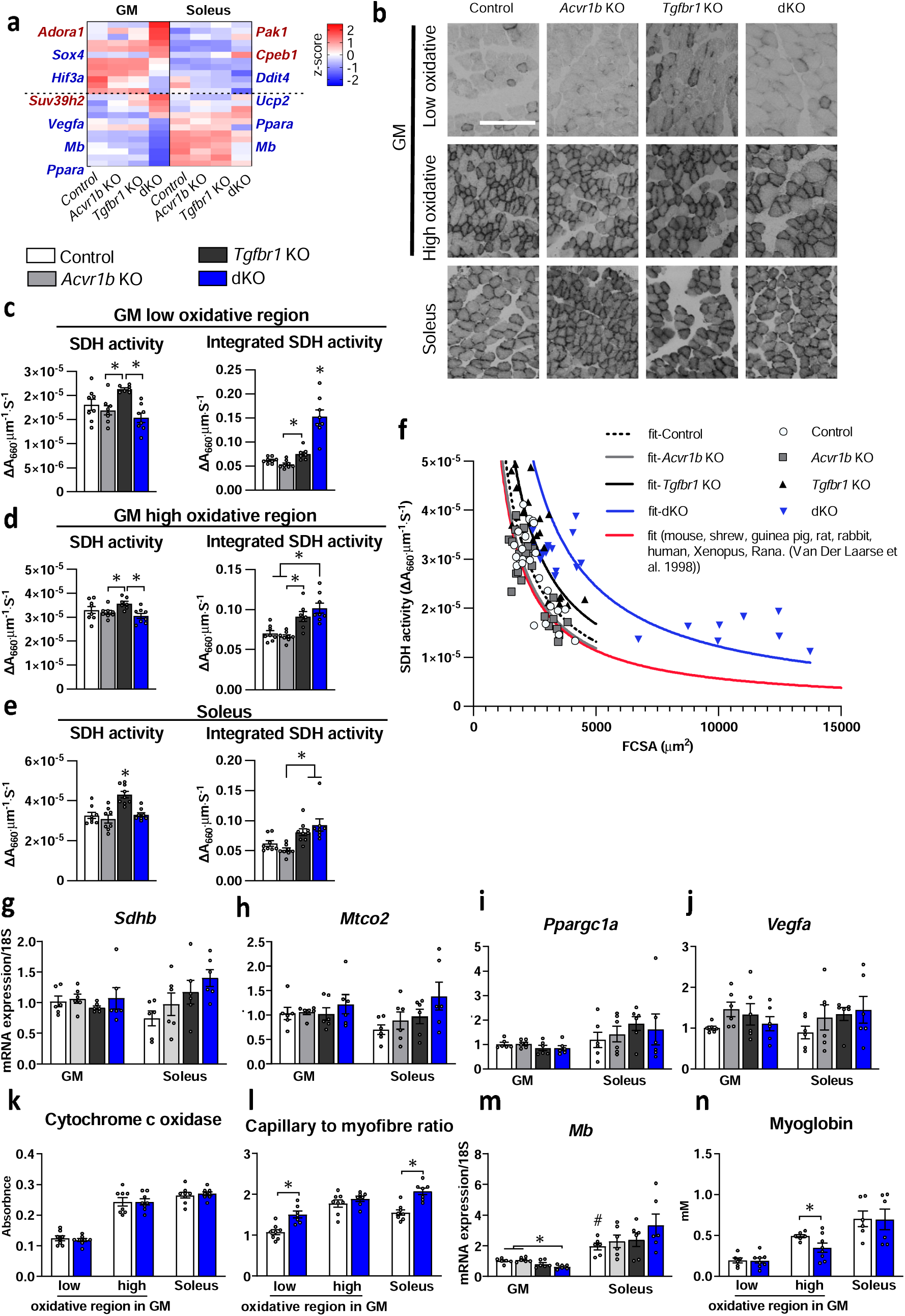
Oxidative metabolism in skeletal muscle shows positive adaption in the absence of TGF-β type I receptors. (**A**) Heatmap of metabolism associated genes (n=4). (**B**) Succinate dehydrogenase (SDH) staining of gastrocnemius and soleus muscles. Scale bar=100μm. Myofibre SDH activity and integrated SDH activity in (**C**) low, (**D**) high oxidative region of gastrocnemius (GM) and (**E**) soleus (n=7-8). (**F**) Relation of SDH and myofibre cross-sectional area (FCSA) of both gastrocnemius and soleus presented by a fitted line which are described by SDH=constant × FCSA^-1^. The constant is calculated as the mean of the products SDH × FCSA. SDH activity of different myofibres at physiological temperatures from different muscles of various species (mouse, shrew, guinea pig, rat, rabbit, human, *Xenopus* and *Rana*) is plotted as a function of FCSA in red. qPCR data for (**G**) *Sdhb*, (**H**) *Mtco2* and (**I**) *Ppargc1a*, (**J**) *Vegfa* (n=6). (**K**) Cytochrome c oxidase activity in myofibres for which the absorbance at 436 nm was not different between control and dKO mice (**L**) Number of capillaries per myofibre (n=7-8). (**M**) Gene expression level of *Mb* (n=6). (**N**) Mean myoglobin protein concentration per myofibre (n=7-8). Two-way ANOVA was used with Bonferroni post hoc tests. Independent t-tests were used to compare data of gastrocnemius and soleus muscles of control mice. Data are shown as mean ±LJSEM. *: *P* < 0.05. A single star indicates a significant difference compared to other groups of the same muscle. #: *P* < 0.05 compared to GM of control mice.

Another determinant of the oxidative capacity is the oxygen supply. Capillary density (mm^-2^) was only less in the high oxidative region in gastrocnemius. Capillary to myofibre ratios in the low oxidative regions in gastrocnemius and in soleus muscles of dKO mice were higher than those in muscles of control mice (**Figure 5. L**, **Supplementary Figure 4. B**, **C**). Genes related to angiogenesis, such as *Angptl2*, *Plau* and *Dll1,* were upregulated in dKO mice as shown by RNA-seq analysis (**Supplementary Data**). Given the improved vascular organization, we next examined the intracellular oxygen transport capacity. Although gene expression levels of *Mb* were reduced in dKO gastrocnemius, myoglobin concentration was only reduced in high oxidative region. Since the high oxidative region comprises less than 40% region of a gastrocnemius (**Figure 5. M**, **N**, **Supplementary Figure. 2)**, this suggests that facilitated intracellular oxygen transport was enhanced in the majority of the myofibres. Taken together, the data imply that despite the substantial hypertrophy, oxidative phosphorylation activity was improved in muscles lacking both TGF-β type I receptors.

## DISCUSSION

Here, we investigated the changes in muscle transcriptome, contractile force and phenotype of both fast, glycolytic and slow, oxidative muscle with myofibre-specific receptor knockout of either individual or both *Acvr1b* and *Tgfbr1*. Our data show that simultaneous myofibre-specific knockout of both TGF-β type I receptors induced the most differentially expressed genes in gastrocnemius muscle, including genes related to muscle growth, contraction, differentiation, filament organization, matrix remodelling, and metabolism, while the transcriptome in soleus of dKO mice was affected to a lesser extent. Moreover, double knockout of both TGF-β type I receptors induced substantial hypertrophy which was accompanied by an increase in muscle force-generating capacity. In contrast, knockout of individual receptors had little effect on the changes of muscle mass and strength. For fast-type muscle in dKO mice, muscle mass was increased by 60%, while maximal isometric contraction force was only increased by 30%, resulting in a 23% reduction in specific force. For slow-type soleus muscle, the magnitude of hypertrophy was substantially lower (23%), the maximal isometric contractile force was increased by 66%, yielding an unaltered specific force. Strikingly, for both fast- and slow-type muscles of dKO mice increased myofibre CSA was accompanied by increased integrated SDH and cytochrome C oxidase activity in muscles, indicating increased oxidative capacity. In addition, capillary to myofibre ratio was increased and myoglobin concentration was unchanged within myofibres in low oxidative region of gastrocnemius and soleus lacking both type I receptors, suggesting enhanced oxygen supply and sustained intracellular oxygen transportation in hypertrophic muscle. Our study reveals a critical role of TGF-β type I receptors in the regulation of muscle type-specific strength, phenotype, metabolism and provides targets and cues for the development of treatment for muscle wasting disorders.

### Fast-type gastrocnemius muscle lacking both TGF-β type I receptors shows the largest number of DEGs and highest increase in muscle mass

Targeting both *Acvr1b* and *Tgfbr1* in fast-type muscles resulted in the largest number of DEGs and sizable muscle hypertrophy compared to control (wild-type) muscles. Simultaneous receptor knockout induced modest DEGs and a lesser extent of hypertrophy in the soleus. Although transcriptomic profiles in both gastrocnemius and soleus of single knockout mice were different from those of wild-type mice, the lack of either *Acvr1b* or *Tgfbr1* induced modest muscle hypertrophy and phenotype alterations. These results imply functional redundancy of ACVR1B and TGFBR1 in skeletal muscle. The larger number of DEGs and muscle hypertrophy in gastrocnemius of dKO mice may contribute to the higher gene expression levels of *Acv1b* and *Tgfbr1* than in soleus. Since gene expression level of *Acvr1b* and *Tgfbr1* in the soleus of control mice was 10% and 40% of that in gastrocnemius of control mice, respectively (**Figure 1. C**, **D**), low expression levels of TGF-β type I receptors likely resulted in a less pronounced effect of gene knockout on muscle transcriptomic profile and phenotype. In addition, most of the DEGs in fast-type muscle of dKO mice were associated with muscle contraction, muscle growth and differentiation, sarcomere organization, matrix remodeling, and metabolism (**Figure 1. K**). These findings indicate that TGF-β signalling plays an important role in several cellular processes and physiological activities in skeletal muscle.

In addition to the largest number of DEGs, double knockout of TGF-β type I receptors induced the largest hypertrophy in fast-type muscle. Previously we have shown that 5 weeks after tamoxifen-induced knockout of both TGF-β type I receptors in skeletal muscle, type IIB myofibre CSA of tibialis anterior muscle was increased by twofold, while type I myofibre CSA was not changed ^40^. Here, three months after inducible knockout of both type I receptors, FCSA was increased in both fast-and slow-type muscles. Interestingly, FCSA in low oxidative region of gastrocnemius muscles which consist predominantly of type IIB myofibre was increased by 3.2-fold in dKO mice (**Figure 2. E**), while FCSA in high oxidative region of gastrocnemius and soleus of dKO mice was increased by 1.5-fold (**Figure 2. G**). These observations are in line with previous observations that fast-type myofibres of human and rodents have a stronger potential to hypertrophy in response to hypertrophic stimuli by resistance exercise ^41,42^ or mechanical overload ^43^. In addition, the increased *Hgf* expression, particularly in gastrocnemius muscles of dKO mice, may have contributed to the substantial muscle hypertrophy in this muscle which exceeded that shown in soleus. HGF induces both protein synthesis signalling in muscle cells and activates MuSCs ^44,45^, the increased expression of *Hgf* in gastrocnemius of dKO mice likely stimulated an increase in protein synthesis rate and may have supported regeneration of myofibres in the outer region (**Figure 4. J**). Since *Hgf* expression was still upregulated after 3 months of dKO mice, it is highly conceivable that HGF was a key factor in the continuous elevated protein synthesis in fast-type muscle.

### Maximum isometric force is increased while specific force is reduced in fast-type muscle in the absence of both TGF-β type I receptors

Despite the substantial muscle hypertrophy induced by double knockout of TGF-β type I receptors in gastrocnemius, muscle maximum isometric force was not proportionally increased, resulting in reduced specific force. In contrast, specific force was not decreased in soleus (**Figure 2. H**, **I**). The faster rate of hypertrophy and the discrepancy in hypertrophy and force-generating capacity in fast-type myofibres raise the question what the mechanism is underling these differences between fast-type and slow-type myofibres and what the explanation is for the differential effects on force-generating capacity. Several factors determine the maximal force-generating capacity of a muscle (e.g., myofibre type composition, cross bridge density, and muscle architecture) ^46^.

The disproportional changes in muscle size and strength in the fast-type myofibres may be explained by two mechanisms. One explanation may be an unbalanced increase in contractile protein content and in systems for excitation-contraction coupling (ECC). Although fast-type myofibres generate higher specific forces than slow-type myofibres ^47,48^, the increase in myofibre CSA occurred concurrently with a reduction in specific force ^49^. Previously it has been shown that supplementation of insulin stimulates hypertrophy of mature single myofibres *in vitro*, while the increase in maximal force remains half of the percentage of increase in CSA when the number of sarcomeres in series is unchanged ^50^. It is conceivable that the production of contractile machinery and functional improvement is delayed compared to the increase in muscle mass ^51^. This observation is in line with the concurrently reduced specific force and myosin content in myostatin-null mice, indicating a reduced number of functional bound cross-bridges ^30^. Here in the current study, the reduced gene expressions of *Mylk2, Myl2, Myh7* and *Myh6* in gastrocnemius of dKO mice suggests a reduced expression of proteins involved in the contractile machinery. The imbalance in contractile and total protein content in myofibres of dKO mice likely contributed to the reduction in specific force of gastrocnemius. In addition, in our transcriptomic analysis, sarcolipin (*Sln*), a negative regulator of sarcoplasmic reticulum ATPase (SERCA), showed a 20-fold increase in gastrocnemius muscles of dKO mice, suggesting an enhanced inhibition of cytosolic calcium reuptake in the sarcoplasmic reticulum. Inhibition of SERCA protein activity disrupts the ECC process, which has been related to increased cytosolic calcium and muscle weakness ^52^. It has been shown that overexpression of *Sln* reduced peak twitch force and tetanic force in soleus muscle ^53^, and also reduced contractility of cardiac myocytes due to the reduced calcium uptake rate of sarcoplasmic reticulum ^54^. The substantially increase in *Sln* expression level in gastrocnemius may explain the reduced specific force. The differences in the fold increase of *Sln* expression between gastrocnemius and soleus (20-fold vs. 2.3-fold) of dKO mice potentially have contributed to the differences in muscle force generating compacity (i.e. specific force).

A second explanation could be the impaired replacement of dysfunctional contractile and cytoskeletal proteins. Our RNA-seq data show that gene expression of *Foxo1* was decreased in gastrocnemius of dKO mice, indicating inhibition of protein degradation (**Supplementary Data**). The inhibition of protein degradation rate may lead to accumulation of dysfunctional myofibrils both quantitively and qualitatively, causing a reduced force production ^55^. These observations are in line with the reduced specific force in EDL muscle of myostatin-null mice, which was accompanied by reduced expression levels of atrogin-1 and ubiquitinated myosin heavy chain^56^. Similarly, muscle-specific knockout of *Foxo1*, *3* and *4* in mice increased myofibre CSA while maximal gastrocnemius contraction force was also reduced ^57^. Skinned myofibre segment experiments are warranted to reveal the contribution of the impact of dysfunctional proteins in the reduction of specific force.

### Lack of both type I receptors reduces eccentric contraction force in gastrocnemius muscles while has minor effects on muscle force-frequency relationship

Susceptibility to eccentric contraction-induced injury was examined for both the soleus and gastrocnemius muscles. For all groups, the relative force exerted during the series of eccentric contractions was less in gastrocnemius muscles of dKO mice rather than in soleus muscles of dKO mice compared to other groups, suggesting that fast, glycolytic muscle is more susceptible to eccentric loading induced strength loss (**Figure 4. A**). This observation was in line with those of other studies showing a larger drop in force in fast, low oxidative EDL muscles compared to slow, high oxidative soleus muscles after eccentric contractions, followed by a higher release of lactase dehydrogenase, indicating more sarcolemmal damage in the fast-type muscle ^26,58^. Type II myofibres are more damaged after eccentric contraction than type I myofibres ^59^. Cytoskeletal elements play critical roles in maintaining myofibre structural integrity. Fibre type relative differences in Z-line composition, the titin isoform stiffness, nebulin size, dystrophin content and length have been argued to contribute to the higher vulnerability to injury and the force loss ^60^. However, we did not observe any differences in the expression of related genes.

Another factor that may have contributed to the enhanced susceptibility to injury is related to the increase in gene expression of *Mybph* (**Figure 3. N**)*. Mybph* encodes a homologue of MYBPC. MYBPH is presumed as a putative skeletal muscle biomarker of amyotrophic lateral sclerosis and signals the onset of disruption of actin-myosin interaction ^61^. Upregulation of *Mybph* is a feature of severe myopathy ^62,63^. The extent to which the elevated *Mybph* expression in dKO muscles contributes to the susceptibility of eccentric contraction-induced strength loss remains to be determined.

In addition, we observed an increase in *Obsl1* expression in both fast- and slow-type muscles of dKO mice (**Figure 3. M**). *Obsl1* encodes for obscurin-like 1 (OBSL1) which is a homologue of obscurin. Both obscurin and OBSL1 bind to myomesin and titin in the M-band to stabilize sarcomere structure ^64^. The increased *Obsl1* expression suggests an adaptation of the cytoskeleton in response to the high forces that was exerted by hypertrophic myofibres. Despite the elevated levels of *Obsl1* in dKO mice, the substantial loss in gastrocnemius muscle strength of dKO suggests that assembly of cytoskeleton proteins was not optimal in fast-type muscle. As the function of OBSL1 is not fully understood yet, the disruption of sarcomeric alignment and actin-myosin function during eccentric contraction in fast-type muscle of dKO mice requires further investigation.

Alternatively, high forces generated by the hypertrophic myofibres may cause high, local sarcomere strains along the sarcolemma, resulting in an increased susceptibility to contraction induced injury. Moreover, as part of the cytoskeleton, actin filaments are linked to the syntrophin complex by dystrophin. In acinar cells, knockout of *Sntb1* has been shown to result in dispersed actin and defective actin assembly ^65^. Here, reduced expression of *Sntb1* in gastrocnemius muscles of dKO mice is suggested to destabilize the sarcomeres. Given that dystrophin-deficient mice are particularly susceptible to damage from eccentric contractions due to the disruption of DGC complex ^66^, it is conceivable that in dKO gastrocnemius the reduced *Sntb1* expression, as one of the DGC elements, may jeopardize cell membrane integrity and increase susceptibility to contraction-induced muscle damage ^67^. As a result, the scar tissue and regenerating myofibres observed in regions of in low oxidative region of dKO gastrocnemius may lead to further muscle injury and force loss after eccentric contraction (**Figure 4. J**). In addition, reduced *Sntb1* expression levels have been shown to be associated with increased myotube diameter *in vitro*, suggesting the regulatory effect of SNTB1 on muscle size ^68^.

This study shows that endomysium and perimysium thickness in muscles lacking both type I receptors were increased in proportion with the increase in myofibre CSA (**Figure 4. H**, **I**). The increase endomysium and perimysium thickness could be an adaptation to the higher forces exerted by hypertrophic myofifbre ^69^ and as such does not explain the enhanced eccentric contraction-induced reduction in force-generating capacity in dKO gastrocnemius muscle.

### Lack of both Acvr1b and Tgfbr1 stimulates hypertrophy and oxidative metabolic capacity

Here we demonstrate that simultaneous myofibre-specific knockout of both TGF-β type I receptors allows for simultaneous increase myofibre size and oxidative capacity. Myofibres that lack both *Tgfbr1* and *Acvr1b* show increased integrated SDH activity compared to myofibres of similar size of control mice and other species. In general, throughout a range of species myofibre size and oxidative metabolism are inversely related ^13,39^, indicating that hypertrophy and an increase in oxidative metabolism are mutually exclusive. Generally, it is a challenge to simultaneously increase muscle size and oxidative capacity cf.^13^. In human, the increase in muscle mass induced by resistance training was blunted by concurrently performing endurance training which increased mitochondrial activity ^70,71^. Our quantitative histochemical assessment of SDH activity provides calibrated histological estimates of myofibre VO_2max_ ^72^. Integrated SDH activity was increased in myofibres of dKO mice. In addition, cytochrome c oxidase activity per myofibre CSA was not changed in hypertrophic myofibres of dKO mice compared to control (**Figure 5. K**). The increased integrated SDH and cytochrome c oxidase activities in myofibres of dKO mice indicate the increased mitochondrial respiratory chain activity. This unique finding demonstrates that the inverse relation between of myofibre size and oxidative metabolism capacity is violated by knocking out both TGF-β type I receptors.

Regarding the regulatory mechanisms underlying the increase in myofibre mitochondrial biosynthesis, the transcriptome analysis suggested that the increased oxidative metabolic capacity was due to the increased transcription of mitochondrial genes per myofibre. Since the amount of total mRNA per milligram muscle and the number of total nuclei per myofibre were not changed (**Supplementary Figure 1. J, Figure 3 H**), the increased mRNA levels per myofibre was due to the increase in transcripts. This suggests that in dKO mice, transcriptional rate of mitochondrial genes per myonucleus was enhanced, which contributes to the increased number of proteins in the respiratory chain and mitochondrial content, leading to increased aerobic metabolic activity in myofibres. In addition, 20-fold increased *Sln* in gastrocnemius of dKO mice may contribute to the enhanced aerobic metabolism. High *Sln* expression level has been shown to increase oxidative metabolism in skeletal muscle ^73^. Over expression of *Sln* triggered mitochondrial biogenesis by raising intracellular calcium concentration which activates CAMKII and PGC-1α ^74^. By uncoupling SERCA, SLN increased energy demand, leading to enhanced mitochondrial biogenesis and ATP production ^75^. These observations indicate that TGF-β type I receptor signalling is likely a key play in the regulation of mitochondrial biosynthesis.

Sufficient oxygen supply is pivotal to maintain mitochondrial ATP production ^76^. We asked whether increased integrated SDH activity in dKO mice was accompanied by increased oxygen supply and uptake. Previous studies have shown that capillary supply to a myofibre is associated with the myofibre CSA ^77^. Local capillary to myofibre ratio showed a positive correlation with myofibre size in human and rates muscles ^78^. We assumed that intracellular oxygen transportation should be increased to meet the oxygen demand within myofibres lacking both *Acvr1b* and *Tgfbr1*. Here, we found an increase in capillary to myofibre ratio in the low oxidative region in gastrocnemius and soleus muscles in the absence of both type I receptors (**Figure 5. N**). This indicates that myofibre-specific knockout of both TGF-β type I receptors stimulates capillarization. In addition to extracellular oxygen supply, intracellular oxygen transportation is also important to facilitate oxygen diffusion for oxidative energy production, which is mainly regulated by myoglobin ^79,80^. Although gene expression levels of *Mb* in gastrocnemius of dKO mice were reduced (**Figure 5. L**), myoglobin concentration within myofibres was unchanged in the low oxidative region of gastrocnemius and soleus (**Figure 5. M**), which was likely caused by enhanced rate of translation. This suggests that increased myoglobin content may have contributed to enhanced intracellular oxygen transport within the superficial region of fast-type muscle and slow-type muscle to prevent hypoxia in the core of myofibre. In addition to oxygen storage and diffusion to the mitochondria within the myocyte, deoxymyoglobin has been shown to have strong nitrite reductase activity and able to generate nitric oxide (NO) ^81^. NO causes vasodilation and increases the blood flow to ensure sufficient oxygen delivery to the myofibre to match its metabolic requirements. Overall, the myofibre-specific lack of both *Acvr1b* and *Tgfbr1* in skeletal muscle increases myofibre CSA, oxidative metabolism, vascularization and facilitates intracellular myofibre oxygen transport simultaneously, suggesting that inhibition of TGF-β type I receptors signalling potentially improves muscle endurance capacity.

### Potential clinical implications of simultaneous knockout of Acvr1b and Tgfbr1

The present study shows three results with clinical relevance. First of all, myofibre-specific lack of both *Acvr1b* and *Tgfbr1* increases myofibre size and oxidative capacity, indicating that potential pharmacological interference with these receptors may improve muscle strength and endurance capacity simultaneoulsy. Secondly, our results indicate that simultaneous receptor knockout positively affects force generation capacity particularly in slow, high oxidative myofibres, as a specific force in soleus muscles is preserved. Increased muscle mass with a less than proportional increase in force-generating capacity may not be beneficial. It is worth investigating whether endurance exercise combined with simultaneous pharmacological inhibition of *Acvr1b* and *Tgfbr1* may enhance force-generating capacity, while preserving or improving specific muscle force. Note that human skeletal muscles do not express type IIB myosin, thus simultaneously blocking *Acvr1b* and *Tgfbr1* in adult human skeletal muscle may resemble the results observed in mouse soleus muscle rather than those in gastrocnemius muscle. Last, lack of *Acvr1b* and *Tgfbr1* in fast, glycolytic muscles reduced eccentric force-generating capacity, which is a potential indication for increased susceptibility to injurious eccentric loading. Therefore, it is noteworthy that in the development of a therapeutic intervention, substantial hypertrophy of glycolytic myofibres should be restrained to minimize the risk for injury.

In conclusion, our results indicate that myofibre-specific long-lasting simultaneous blocking of *Acvr1b* and *Tgfbr1* in mice skeletal muscles results in muscle transcriptional changes, including genes involved in muscle growth, contraction, differentiation, filament organization, matrix remodelling, and metabolism. Knockout of both type I receptors differentially increases muscle mass and improves force-generating capacity in fast, glycolytic gastrocnemius and slow, high oxidative soleus muscles. Combined inhibition of both receptors in fast-type muscle induces a larger increase in muscle mass than that in force-generating capacity, while muscle mass in slow-type muscle is increased to a lower extent, but is in proportion with the increase in force-generating capacity. Substantial hypertrophy in fast-type muscles increases susceptibility to eccentric contraction induced injury. Moreover, combined myofibre specific knockout of both *Tgfbr1* and *Acvr1b* increases SDH activity in proportion with the increased myofibre size. Although myofbre size and oxidative capacity are usually inversely related, simultaneous inhibition of TGF-β type I receptors seems to be a beneficial strategy to alleviate muscle pathology that may outperform other strategies, as interference with these receptors may improve muscle strength and aerobic metabolism capacity simultaneously.

## MARTERIALS AND METHODS

### Treatment of transgenic mice

The HSA-MCM mouse line originated from McCarthy lab ^82^ was obtained from Jackson Laboratory (Bar Harbor, ME, USA (stock number # 025750)). Mouse line *Acvr1b^fl/fl^*^83^ was provided by Philippe Bertolino (Cancer Research Center of Lyon, French Institute of Health and Medical Research) and the *Tgfbr1* KO mouse line ^84^ was provided by Peter ten Dijke (Leiden University Medical Center). All animals were of a C57BL/6 background. Mouse lines were crossbred in-house to obtain HSA-MCM:*Acvr1b^fl/fl^*, HSA-MCM:*Tgfbr1^fl/fl^* and HSA-MCM: *Acvr1b^fl/fl^:Tgfbr1^fl/fl^*mouse lines (further be referred to *Acvr1b* KO, *Tgfbr1* KO and dKO mice). Mouse lines were crossbred homozygously for the targeted loxP sites and heterozygously for inducible Cre recombinase enzyme. Food (Teklad, Envigo, Horst, The Netherlands) and water were available ad libitum. All experiments were performed according to the national guidelines approved by the Central Committee for Animal Experiments (CCD) (AVD112002017862) and the Institute of Animal Welfare (IvD) of the Vrije Universiteit Amsterdam. Genotyping for the *HSA-MCM*, *Acvr1b^fl/fl^* and *Tgfbr1^fl/fl^* genes was performed by using DNA isolated from ear notches. PCR was performed according to the corresponding genotyping protocol to validate the effect of gene knockout. The following primer sequences were used for genotyping: *HSA-MCM* gene: forward, 5′- GCATGGTGGAGATCTTTGA-3′ ^85^, reverse, 5′- CGACCGGCAAACGGACAGAAGC-’3 ^82^, *Acvr1b^fl/fl^*gene: *In4*, 5’- CAGTGGTTAAGAACACTGGC-3’, *In5*, 5’- GTAGTGTTATGTGTTATTGCC -3’, *In6*, 5’GAGCAAGAGTTTCTCTATGTAG-3’ ^83^and *Tgfbr1^fl/fl^* gene: forward, 5’- CCTGCAGTAAACTTGGAATAAGAAG-’3, reverse, 5’- GACCATCAGCTGTCAGTACCC-3’ (Protocol 19216: Standard PCR Assay - Tgfbr1<tm1.1Karl>, Jackson Laboratory, Bar Harbor, ME,USA) ^84^. At an age of six weeks old, male mice were injected intraperitoneally with 100mg/kg/day TMX (10mg/ml, Sigma-Aldrich, T5648) for five consecutive days to activate Cre recombinase enzyme.

### RNA-seq and data analysis

RNA from both left gastrocnemius medialis (gastrocnemius) and soleus muscles of four animals was extracted for RNA-seq by using TRIzol (Thermo Fisher Scientific). Ribopure Link (Thermo Fisher Scientific) and DNAse I (Invitrogen) were used to isolate RNA. RNA purity was checked by 2200 TapeStation (Agilent Technologies), RIN values were all above 7.7. RNA concentration was measured by fluorometric quantification QbuitFluorometer (Life Technologies). RNA-seq library was prepared by enriching mRNA with KAPA mRNA Hyperprep (Roche Sequencing Solutions, Indianapolis, IN, USA). A total of 32 libraries were pooled and loaded on 4 lanes of Hiseq 4000 (Illumina, San Diego, CA, USA) that generated a single read of 50-base pairs per sample. On average, 43 million reads were produced per lane. Sequence quality was checked by FASTQC (version 0.11.9) that all sequences past Q value 30. Cutadapt was used for adapter trimming. Sequence reads were mapped to the mouse reference genome (GRCm38.p6, https://www.ncbi.nlm.nih.gov/assembly/GCF_000001635.26/) by HISAT2 (version 1.33). Mapped sequences were converted to normalized read counts using featureCounts (version 1.5.0-p3) ^86^ and Ensembl GTF annotation. DESeq2 was used to analyse the DEGs on R2 platform (https://hgserver2.amc.nl/). Genes with relative expression of Benjamini-Hochberg-adjusted *P* < 0.05 and fold change ≥1.5 were considered differentially expressed. DEGs in gastrocnemius of dKO mice were used to construct heatmaps on R2 platform. GO analysis of biological process was performed on DAVID 6.8 ^87^.

### qPCR assay

Six biologically independent samples were used for qPCR using the Quant Studio 3 real time PCR (Applied Biosystems, Foster City, CA, USA) and SYBR Green master mix (Applied Biosystems, A25742, Foster City, CA, USA). Relative expression levels were normalized to expression levels of 18S mRNA. The primer sets are shown in **Supplementary Table 5**.

### In situ contractile force measurement

Contractile force characteristics of both gastrocnemius and soleus muscles were measured three months after TMX injections. Mice were given analgesic, buprenorphine (Temgesic, 0.3 mg/ml, 0.1mg/kg), and anesthetised with 0.5-2% isoflurane (TEVA Pharmachemie BV, Haarlem) oxygen/air 0.2L/min. Right gastrocnemius medialis and soleus muscles were exposed that the proximal tendon was solely connected to the hind limb while the distal tendon was connected to a force transducer, leaving the innervation and blood supply intact. Muscles were kept humid during the whole assay. An electrical cuff was used to stimulate the gastrocnemius or soleus branch of the tibial nerve. After determining the amplitude of the electrical current for optimal muscle stimulation in mA (DS3 Isolated Current Stimulator, Digitimer), all experiments were performed at this amplitude. The length-force relation was determined by inducing tetanic isometric contractions (300ms, at 150Hz) at increasing muscle lengths, ranging from near active slack length until approximately 1 mm over optimal muscle length (l_o_) (1 or 0.5-mm increments). Here, l_o_ is the muscle length at which the muscle attained the highest active force (P_o_), i.e. the total force minus the passive force generated at that length. Each tetanic contraction was preceded and followed by two twitches with a 300ms interval between each twitch or tetanus. Between two subsequent tetanic contractions, muscles were kept at rest at slack length for 2 minutes to allow metabolical recovery. To determine passive and active muscle lengths during contractions, videos of contractions were analysed in Tracker software version 5.0.7 (obtained September 2019 via https://www.physlets.org/tracker/). Force-frequency relations were determined by inducing tetanic isometric contractions at approximately l_o_ at increasing stimulation frequencies. Subsequently, to test muscle susceptibility to damage, fifteen consecutive eccentric contractions were performed. Muscles were stimulated at 95% of their optimal length at 150Hz for 100ms and stretched to 116% of their optimal length (l_o_) with a speed of 0.75l_o_ (ms^-1^) during stimulation. Total stimulation duration was 500ms. After this, mice were euthanized with 0.1ml Euthasol injected into the heart and muscles were quickly dissected and weighed. Muscles were snap frozen in liquid nitrogen and stored at -80°C.

### Immunofluorescence and histological analysis

Cryosections of 10 μm were cut from both gastrocnemius and soleus muscles (n=6-8). Samples were subjected to immunofluorescence staining using antibodies against type I (BAD-5-s) (1 µg/mL), IIA (SC-71) (10 µg/mL), IIX (6H1) (1 µg/mL) and IIB (BF-F3) (1 µg/mL) (DSHB, Iowa City, IA, USA) myosin heavy chain were used as the primary antibodies. Goat anti-Mouse Alexa Fluor® 647 IgG2b (5 µg/mL), Goat anti-Mouse Alexa Fluor® 488 IgG1 (5 µg/mL), Goat anti-Mouse Alexa Fluor® 647 IgM (5 µg/mL), Goat anti-Mouse Alexa Fluor® 488 IgM (5 µg/mL) (Thermo Scientific, Waltham, MA, USA) were used as secondary antibodies. Sections were stained by wheat germ agglutinin (WGA) (1:50) (Fisher Scientific, USA) before being mounted with mounting medium containing 4‘,6-diamidino-2-phenylindole (DAPI) (Brunschwig, H1500, Amsterdam, The Netherlands). Images of all immunofluorescence assays were captured using a fluorescent microscope (Zeiss Axiovert 200M, Hyland Scientific, Stanwood, WA, USA) with a PCO SensiCam camera (PCO, Kelheim, Germany) using the program Slidebook 5.0 (Intelligent Imaging Innovations, Göttingen, Germany).

### Histochemistry staining

Succinate dehydrogenase activity was quantified according to the method reported by Van der Laarse, *et al* ^88^. Images were captured by a Leica DMRB microscope (Wetzlar, Germany) with calibrated grey filters and a CCD camera (Sony XC77CE, Towada, Japan) connected to a LG-3 frame grabber (Scion, Frederick, MD, United States). The absorbances of the SDH-reaction product in the sections were determined at 660 nm using a calibrated microdensitometer and ImageJ ^89^. SDH activity (ΔA_660_ · µm^-1^ ·s^-1^) was measured by the rate of absorbance per section thickness per second [ΔA_660_/(10 µm · 600s)) at 10× magnification. The integrated SDH activity (ΔA_660_· µm ·s^-1^) was defined as SDH activity × myofibre cross-sectional area (FCSA). The curve of fitted SDH is calculated as an average value of constant SDH × CSA ^172^. The value of constant SDH was calculated as the average of integrated SDH activity of each myofibre. Connective tissue was stained by Sirius red staining. Sections were fixed in bouin and stained by 1% Sirius Red ^90,91^. Capillaries were stained by biotinylated lectin (Griffonia simplicifolia) (Vector Laboratories) ^32^. Calibrated myoglobin staining were performed by the method reported by Van der Laarse, *et al* ^92^. Cytochrome c oxidase activity was assessed as previously described by Old, *et al* ^93^.

### RNAscope assay

RNA in situ hybridization (ISH) was performed by using RNAscope (ACDBio) assay, multiplex fluorescent reagent kit (Cat No. 3423280) and following the manufacture’s protocols. Fresh-frozen cross-sections of both gastrocnemius and soleus were used for staining. The probes were RNAscope™ Probe-Mm-Acvr1b (Cat No. 429271), RNAscope™ Probe-Mm-Tgfbr1 (Cat No. 406201), positive control probe (Cat No. 320881) and negative control probe (Cat No. 320871). Images were captured by using a fluorescent microscope (Zeiss Axiovert 200M, Hyland Scientific, Stanwood, WA, USA). Background fluorescent staining was removed by using a negative staining. Images were deconvoluted by Huygens Professional 23.04. Numbers of *Acvr1b* and *Tgfbr1* transcripts within a myofibre were counted from 10 myofibres per muscle by using ImageJ 1.54 (intensity threshold for *Tgfbr1*: 86-65535, *Acvr1b*: 2-65535; size: 4-500 pixels).

### Statistical analysis

Data obtained from physiological measurements were analysed in MATLAB 2018a (MathWorks, Natick, MA, USA). Graphs were made in Prism version 8 (GraphPad Software, San Diego, CA, USA). All data were presented as mean ± standard error of the mean (SEM). Statistical analysis was performed in SPSS version 26 (IBM, Amsterdam, The Netherlands). Statistical significance for multiple comparisons was determined by two-way analysis of variance (ANOVA) or three-way mixed model ANOVA with post-hoc Bonferroni corrections, and *Tgfbr1* and *Acvr1b* were considered between-group independent factors. Significance was set at *P* < 0.05. Either frequency or contraction number were considered as within subject repeated factor. Assumptions for normality (Shapiro-Wilk, *P*=0.05) and equality of variance (Levene’s test, *P*=0.01) were tested. For a three-way repeated measures ANOVA, data sets that violated these assumptions were excluded for statistical analysis. For two-way ANOVA, if assumptions were violated even after transformation, a Kruskall-Wallis nonparametric test was performed. For force-frequency relation, significance was tested at 5, 80, 100 and 125 Hz. For the eccentric contractions, two-way ANOVA was performed for contraction numbers 1, 4, 7, 10 and 14 in gastrocnemius and numbers 1, 6, 7, and 13 in soleus. Other data points were analysed using Kruskall-Wallis analyses because of violation of normality or equal variance.

## ACKNOWLEDGEMENTS

We gratefully acknowledge Philippe Bertolino (Cancer Research Center of Lyon, French Institute of Health and Medical Research) for providing mouse line *Acvr1b^fl/fl^* and Peter ten Dijke (Leiden University Medical Center) for providing the *Tgfbr1^fl/fl^* mouse line. We thank animal caretakers of the Universitair Proefdier Centrum of the Vrije Universiteit Amsterdam, our colleague Wendy Noort, students Elke Schmitz and Bijee Visbeek for their contribution to data analysis. Funding: Prinses Beatrix Spierfonds, grant number W.OR14-17 from MMG Hillege, WMH Hoogaars, RT Jaspers; China Scholarship Council personal grant form A Shi (CSC grant number: 201808440351).

---

**Supplementary Figure 1.**
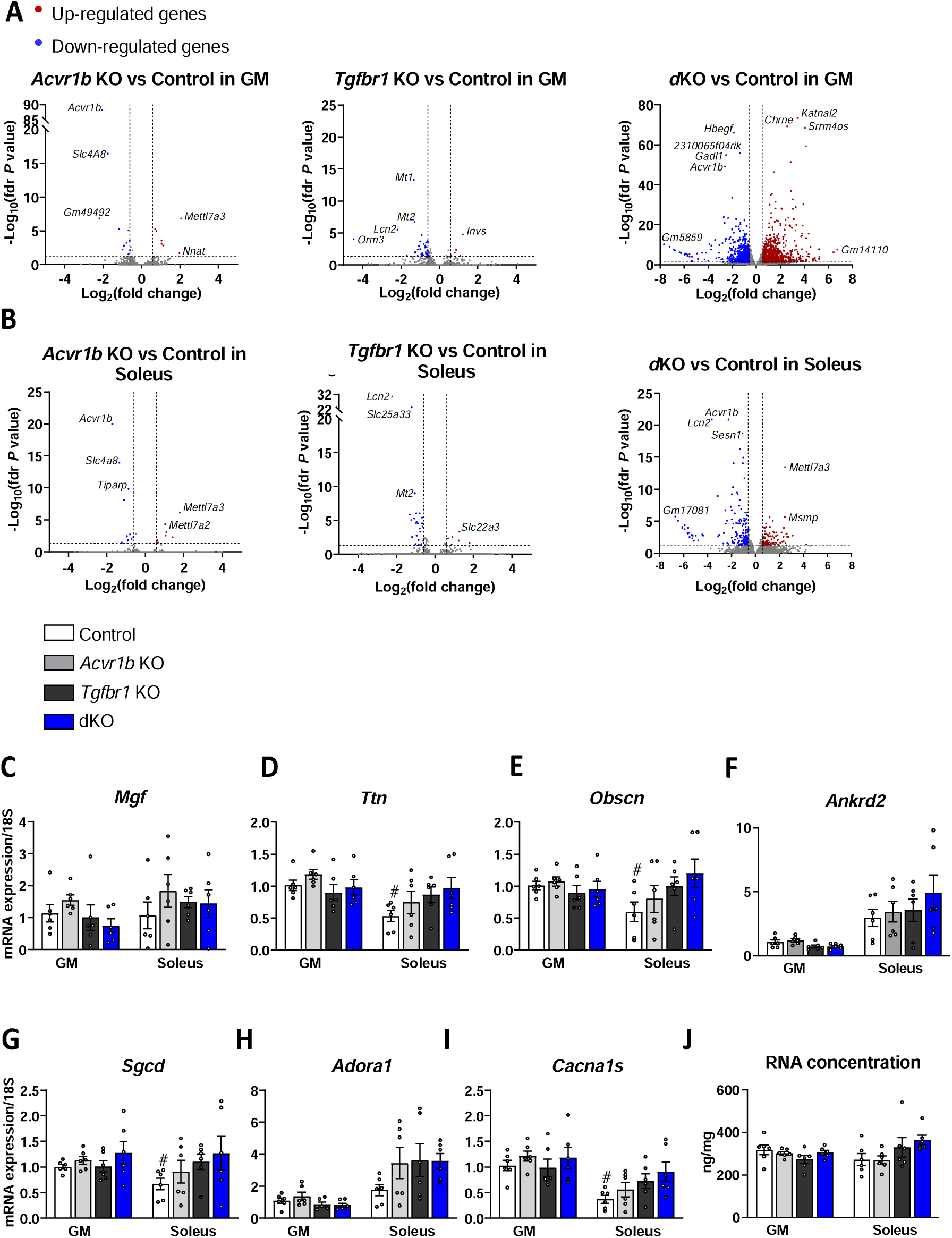
Comprehensive analysis of gene expression by RNA-seq and gene expression analysis by qPCR. (**A**) Volcano plot of differentially expressed genes (DEGs) in gastrocnemius (GM) for each group; there were 8, 7 and 1067 upregulated genes in *Acvr1b* KO, *Tgfbr1* KO and dKO mice respectively; 13, 70 and 744 downregulated genes in *Acvr1b* KO, *Tgfbr1* KO and dKO mice respectively. (**B**) Volcano plot of DEGs in soleus muscles for each group; there were 8, 8 and 85 upregulated genes in *Acvr1b* KO, *Tgfbr1* KO and dKO mice respectively; 13, 46 and 196 downregulated genes in *Acvr1b* KO, *Tgfbr1* KO and dKO mice respectively. Vertical dash lines indicate Log_2_ (fold change) when fold change is ± 1.5. Horizontal dash line indicates -Log_10_ (fdr *P* value) when fdr *P* value is 0.05. DEGs: fdr *P* < 0.05, fold change > 1.5. Gene expression levels of (**C**) *Mgf*, (**D**) *Ttn*, (**E**) *Obscn*, **(F**) *Ankrd2*, (**G**) *Sgcd*, (**H**) *Adora1* and (**I**) *Cacna1s*. (**J**) Total amount of RNA per microgram muscle tissue was not different between groups in either gastrocnemius or soleus muscles. Two-way ANOVA with Bonferroni post hoc tests were performed to analyse data. Independent t-tests were used to compare qPCR data of gastrocnemius and soleus muscles of control groups. Data are shown as mean ±LJSEM. #: *P* < 0.05 compared to GM of control mice.

**Supplementary Figure 2.**
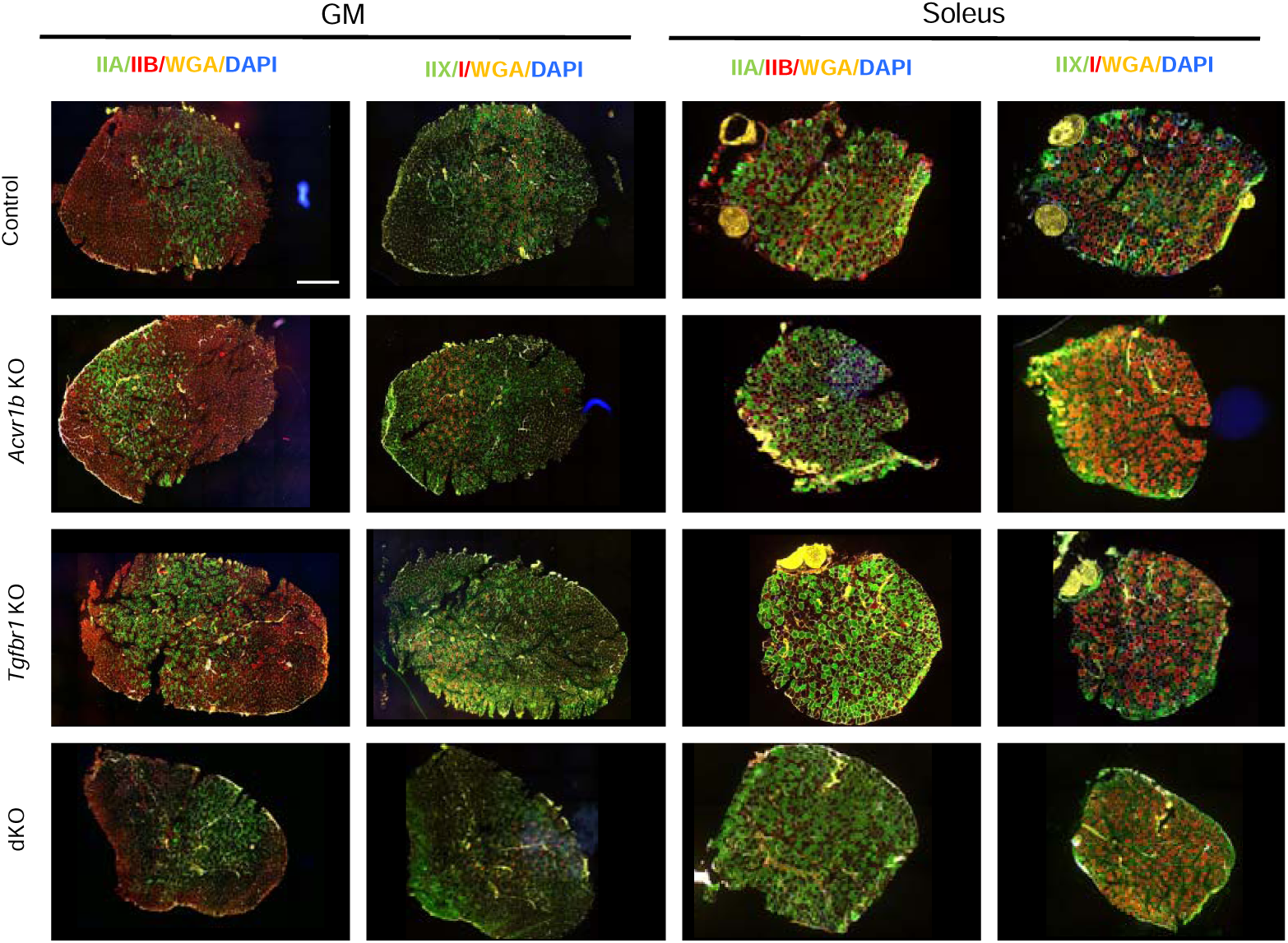
Myofibre type distribution in gastrocnemius and soleus muscles. Immunofluorescence staining of type IIB (red), IIA (green), IIX (green), and I (red) myosin heavy chain in gastrocnemius and soleus muscles. Yellow: wheat germ agglutinin (WGA), blue: DAPI for nuclei. Scale bar=1mm

**Supplementary Figure 3.**
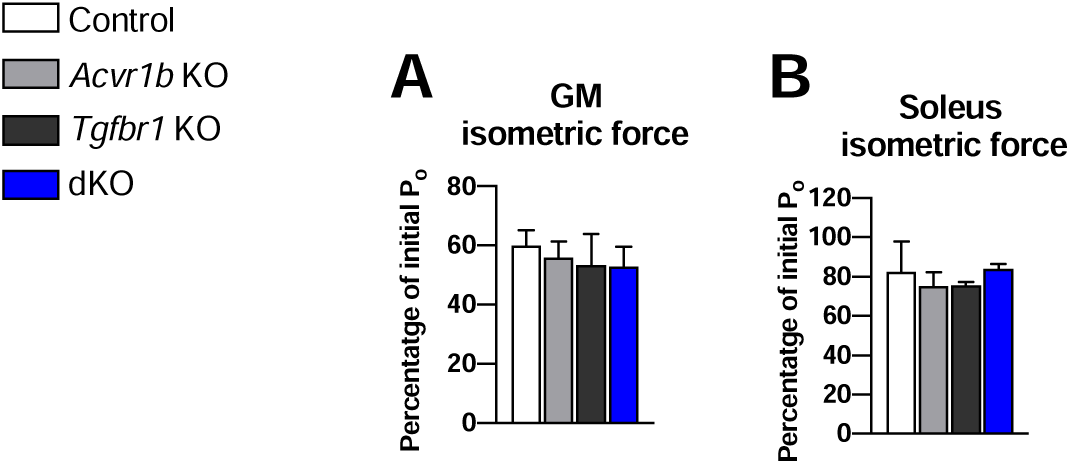
Maximal isometric contraction force (P_o_) after 15 consecutive eccentric contractions. Percentage of P_o_ post 15 consecutive eccentric contractions was reduced compared to P_o_ of pre-contraction. Two-way ANOVA with Bonferroni post hoc tests were performed to analyse data. Data are shown as mean ±LJSEM.

**Supplementary Figure 4.**
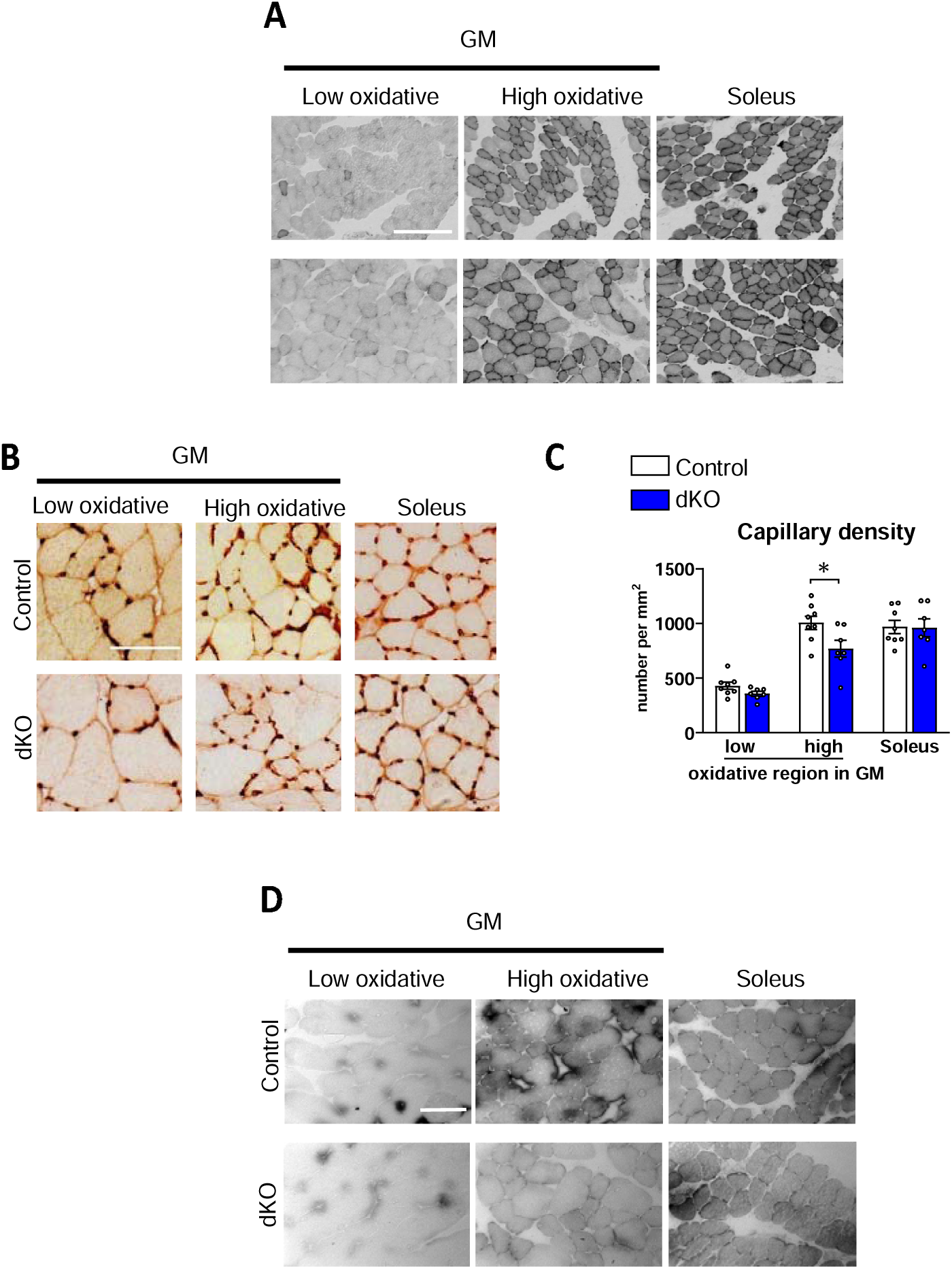
Intracellular oxygen supply and capillary distribution were not diminished in both gastrocnemius (GM) and soleus muscles in dKO mice. (**A**) cytochrome c oxidase, (**B**) capillary and (**D**) myoglobin stainings of gastrocnemius and soleus muscles. For capillary and myoglobin staining, length of scale bars are 100μm. For cytochrome c oxidase staining, length of scale bar is 250μm. (**C**) Capillary density per view was quantified. Independent t-tests were used to compare differences between control and dKO mice. Data are shown as mean ±LJSEM. *: *P* < 0.05.

**Supplementary Figure 5.**
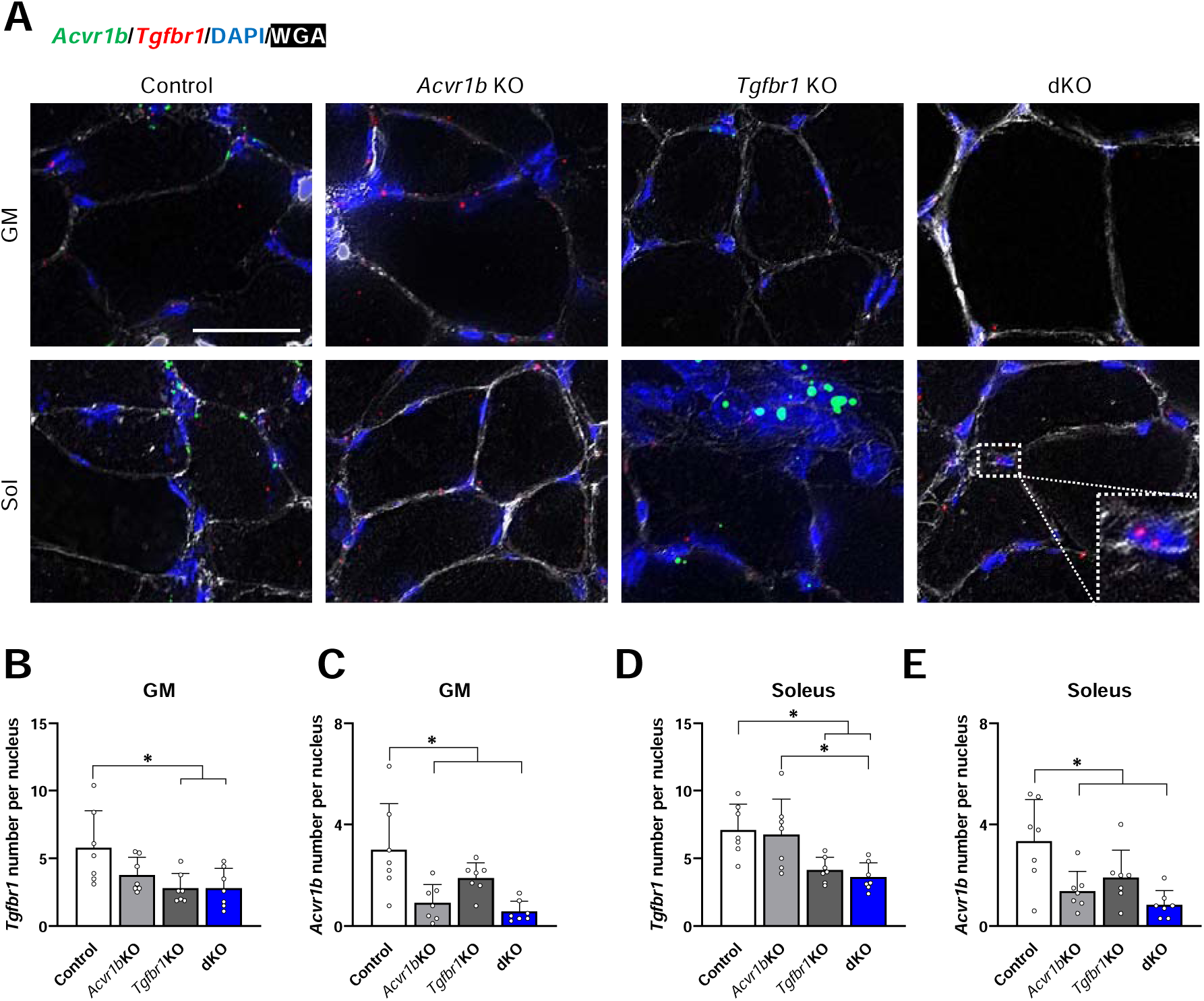
Number of *Acvr1b* and *Tgfbr1* mRNA was decreased per myofibre cross-sectional area (FCSA) in both gastrocnemius (GM) and soleus muscles of corresponding gene knockout mice. (**A**) *Acvr1b* (green) and *Tgfbr1* (red) mRNA was detected by RNA scope assay. White: wheat germ agglutinin (WGA), blue: DAPI for nuclei. Gene expression was found in interstitial cells (dash square). Scale bar=25μm. mRNA number of *Acvr1b* per myofibre CSA in (**B**) gastrocnemius and (**D**) soleus muscles. mRNA number of *Tgfbr1* per myofibre CSA in (**C**) gastrocnemius and (**E**) soleus muscles (n=3). Kruskal-Wallis analysis was used to perform the multiple comparison. Data are shown as mean ±LJSEM. *: *P* < 0.05.

**Supplementary Table 1.** Mouse body and of muscle mass (g)

|  | Control | *Acvr1b* KO | *Tgfbr1* KO | dKO |
| --- | --- | --- | --- | --- |
| Body mass | 34.3±1.2 | 34.5±0.6 | 34.0±0.9 | 39.7±1.1 *† |
| Soleus | 16.6±0.9 | 18.2±0.5 | 19.3±0.9 | 20.6±0.8* |
| Gastrocnemius medialis | 75.2±3.3 | 78.6±3.1 | 84.8±3.1 | 123.7±4.2*#† |
| Gastrocnemius lateralis | 116.6±4.2 | 120.8±3.6 | 126.2±3.8 | 186.6±3.7*#† |
| Plantaris | 25.4±0.4 | 28.1±1.2 | 30.3±1.4 | 40.2±1.4*#† |
| Tibialis anterior | 61.5±1.4 | 65.8±1.4 | 67.0±3.1 | 132.6±4.3*#† |
| Extensor digitorum longus | 14.0±0.2 | 15.3±0.4 | 16.7±0.5* | 26.1±1.1*#† |

N =7-8. \*Significantly different from control, # Significantly different from *Acvr1b* KO, †Significantly different from *Tgfbr1* KO

**Supplementary Table 2.**
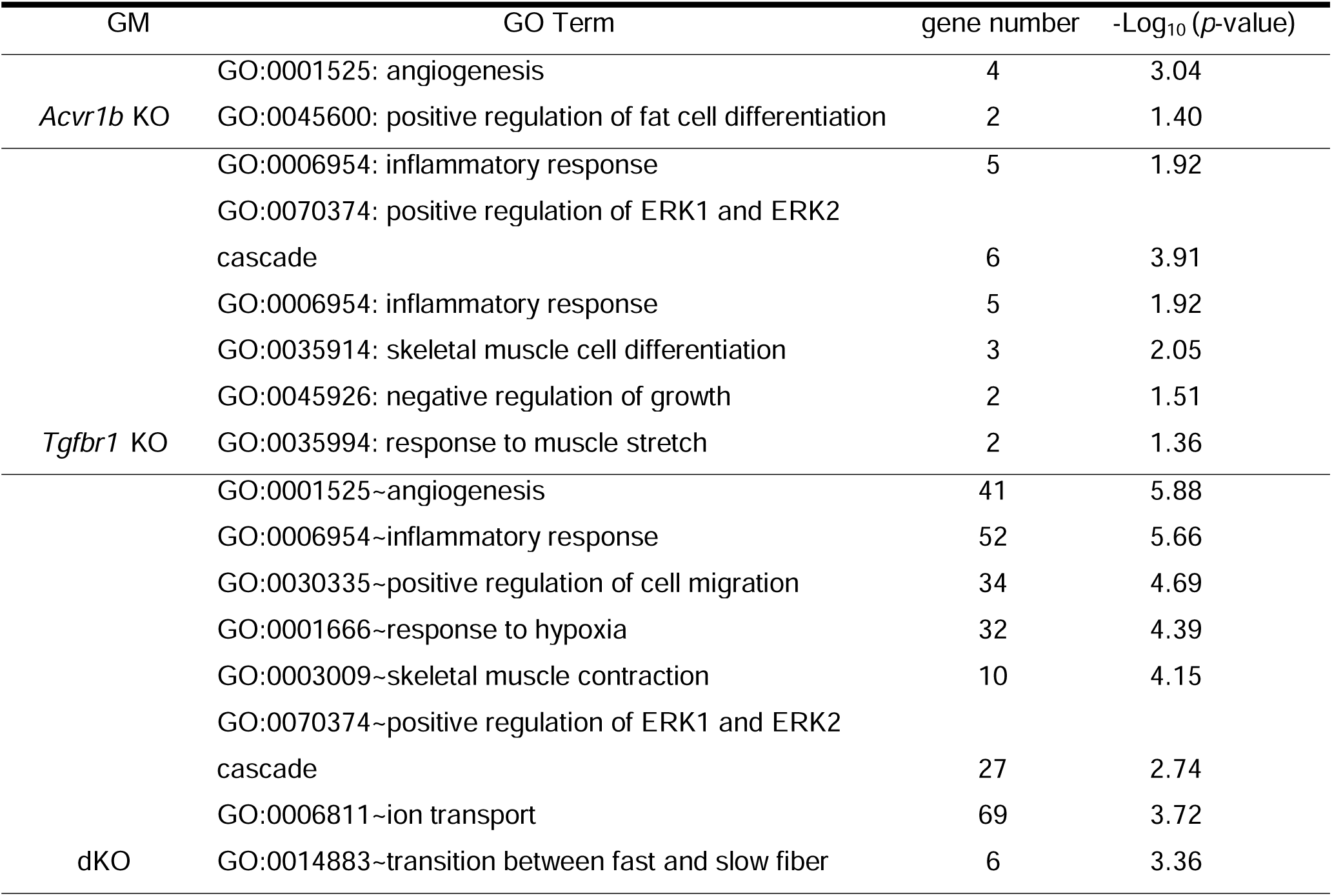
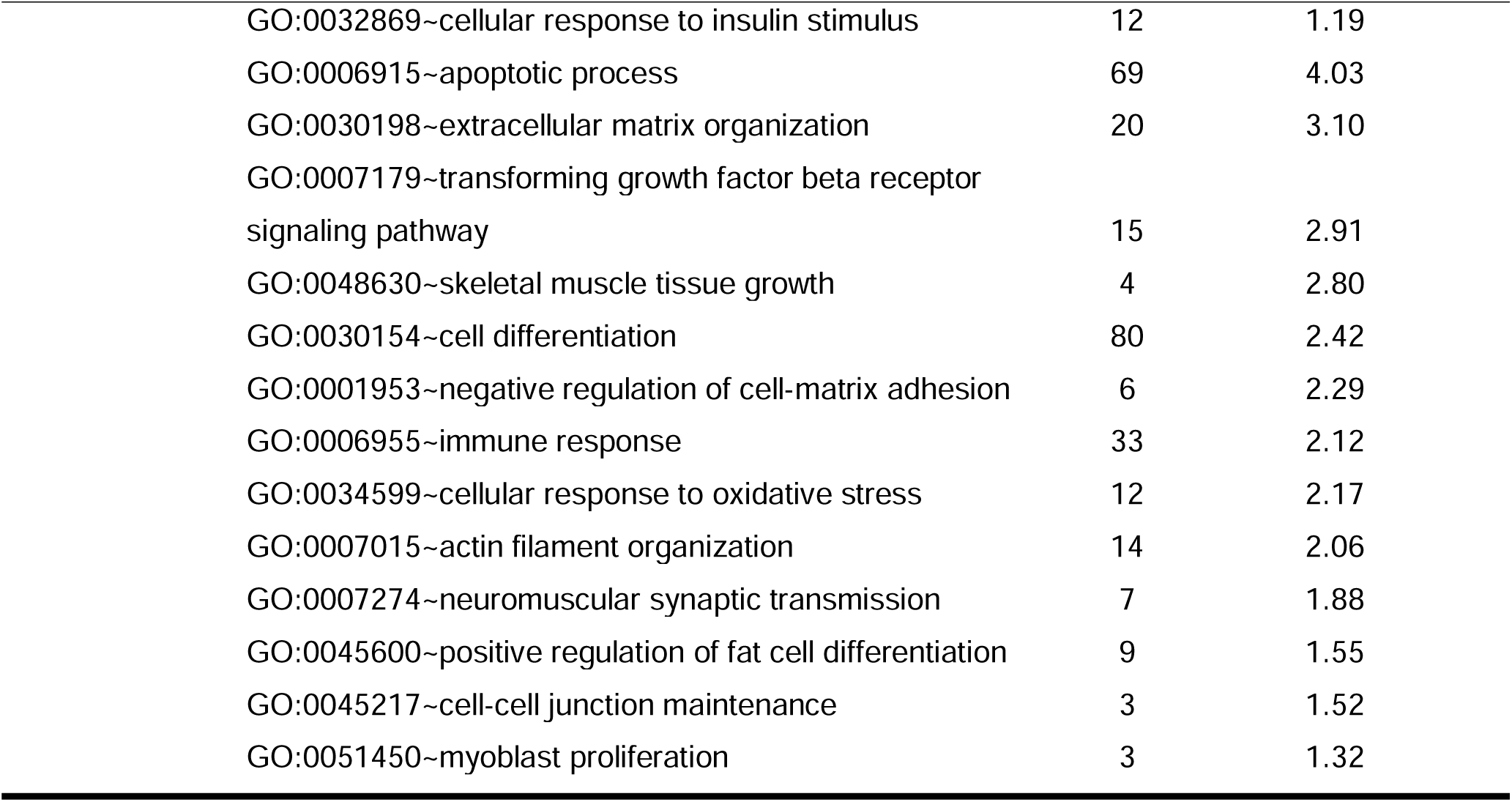
Gene ontology enrichment of gastrocnemius.

**Supplementary Table 3.** Gene ontology enrichment of soleus.

| Sol | GO Term | gene number | -Log<sub>10</sub>(p- value) |
| --- | --- | --- | --- |
| *Acvr1b* KO | GO:0032869~cellular response to insulin stimulus | 2 | 1.37 |
|  | GO:0051443~positive regulation of ubiquitin-protein transferase activity | 2 | 2.03 |
|  | GO:0006468~protein phosphorylation | 3 | 1.50 |
| *Tgfbr1* KO | GO:0032869~cellular response to insulin stimulus | 4 | 3.22 |
|  | GO:0006915~apoptotic process | 7 | 3.22 |
| dKO | GO:0001525~angiogenesis | 8 | 1.92 |
|  | GO:0001666~response to hypoxia | 11 | 4.51 |
|  | GO:0032869~cellular response to insulin stimulus | 6 | 2.67 |
|  | GO:0006915~apoptotic process | 15 | 2.63 |
|  | GO:0050873~brown fat cell differentiation | 4 | 2.30 |
|  | GO:0001568~blood vessel development | 5 | 2.21 |
|  | GO:0006468~protein phosphorylation | 13 | 1.79 |

**Supplementary Table 4.** Physiological properties of gastrocnemius and soleus muscles.

|  | Control | *Acvr1b* KO | *Tgfbr1* KO | dKO |
| --- | --- | --- | --- | --- |
| Gastrocnemius |  |  |  |  |
| P<sub>o</sub> (mN) | 1669.65±105.47 | 1791.03±68.69 | 1743.61±121.34 | 2123.46±142.71 |
| P<sub>t</sub> (mN) | 268.88±25.29 | 326.71±22.80 | 367.62±39.98 | 363.80±44.23 |
| Twitch/tetanus ratio | 0.16±0.01 | 0.18±0.02 | 0.21±0.02 | 0.17±0.01 |
| TTPT (ms) | 26.18±0.18 | 27.30±0.70 | 27.61±0.55 | 26.47±0.16 |
| ½ RT (ms) | 11.68±0.17 | 12.54±0.47 | 12.61±0.36 | 11.97±0.15 |
| dF/dt (mN/ms) | 55.66±3.10 | 53.71±3.11 | 55.13±4.46 | 66.74±5.48 |
| (dF/dt)/P<sub>o</sub> (ms<sup>-1</sup>) | 3.36E-02±1.23E-03 | 3.00E-02±1.43E-03 | 3.17E-02±1.41E-03 | 3.12E-02±7.75E-04 |
| Soleus |  |  |  |  |
| P<sub>o</sub> (mN) | 240.96±21.28 | 281.08±13.69 | 297.04±13.76 | 399.02±10.55 *# |
| P<sub>t</sub> (mN) | 33.24±4.03 | 37.56±3.85 | 46.64±3.36 | 51.60±3.23 * |
| Twitch/tetanus ratio | 0.15±0.02 | 0.14±0.01 | 0.16±0.01 | 0.13±0.01 |
| TTPT (ms) | 34.29±4.17 | 33.19±1.34 | 31.36±0.72 | 30.96±0.46 |
| ½ RT (ms) | 22.61±5.65 | 22.63±2.95 | 18.97±1.13 | 17.67±0.60 |
| dF/dt (mN/ms) | 4.29±0.49 | 5.23±0.37 | 6.04±0.49 | 7.59±0.35*# |
| (dF/dt)/P<sub>o</sub> (ms<sup>-1</sup>) | 1.83E-02±1.49E-03 | 1.89E-02±1.10E-03 | 2.05E-02±1.18E-03 | 1.96E-02±7.47E-04 |

N =7-8. P<sub>o</sub>= maximal tetanic force, P<sub>t</sub>= maximal twitch force; TTPT = time to peak twitch tension; ½ RT = twitch half-relaxation time; dF/dt = maximum rise in tension; dF/dt/P<sub>o</sub>= relative maximum rise in tension.
\*Significantly different from Control, # Significantly different from *Acvr1b* KO, †Significantly different from *Tgfbr1* KO.

**Supplementary Table 5.**
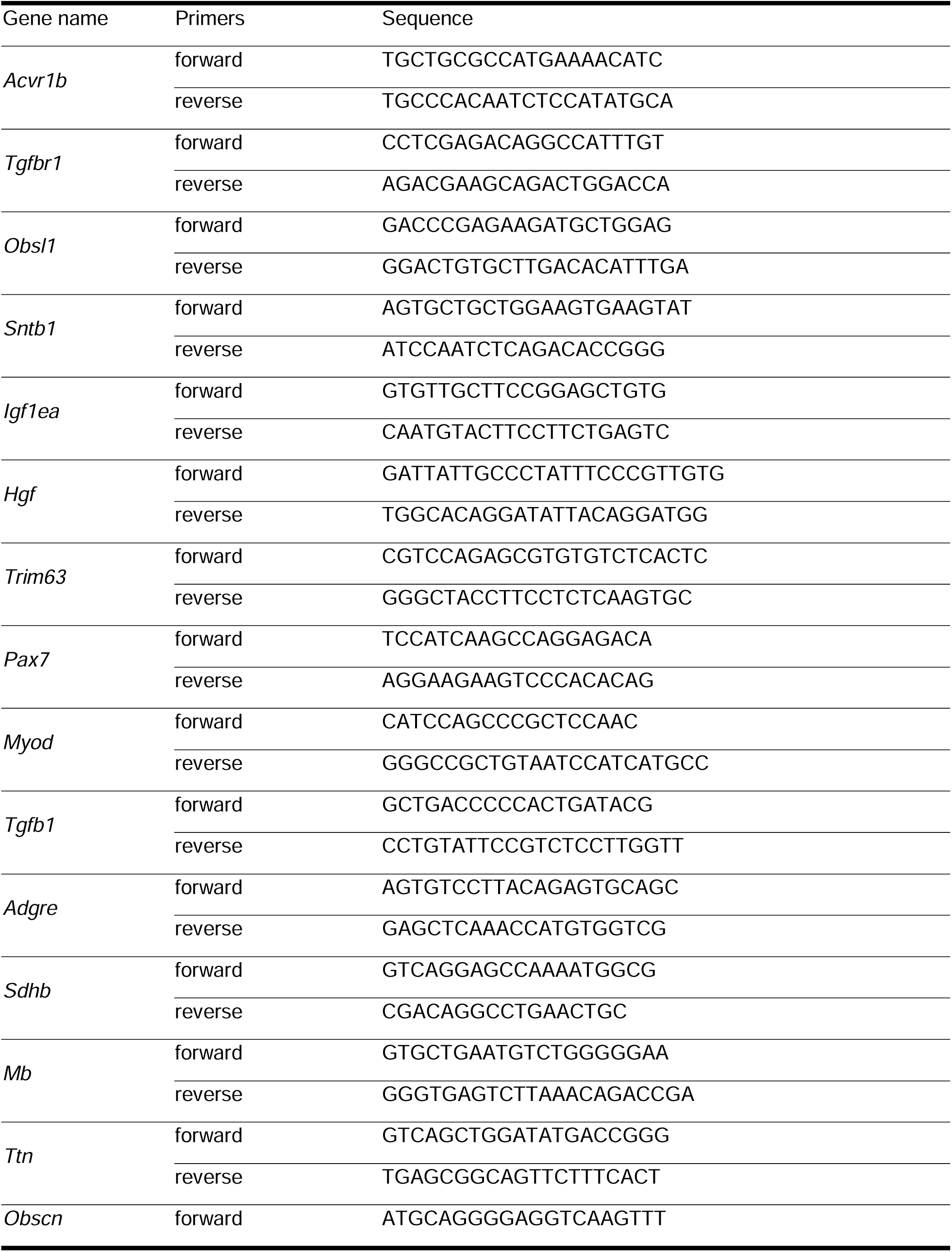
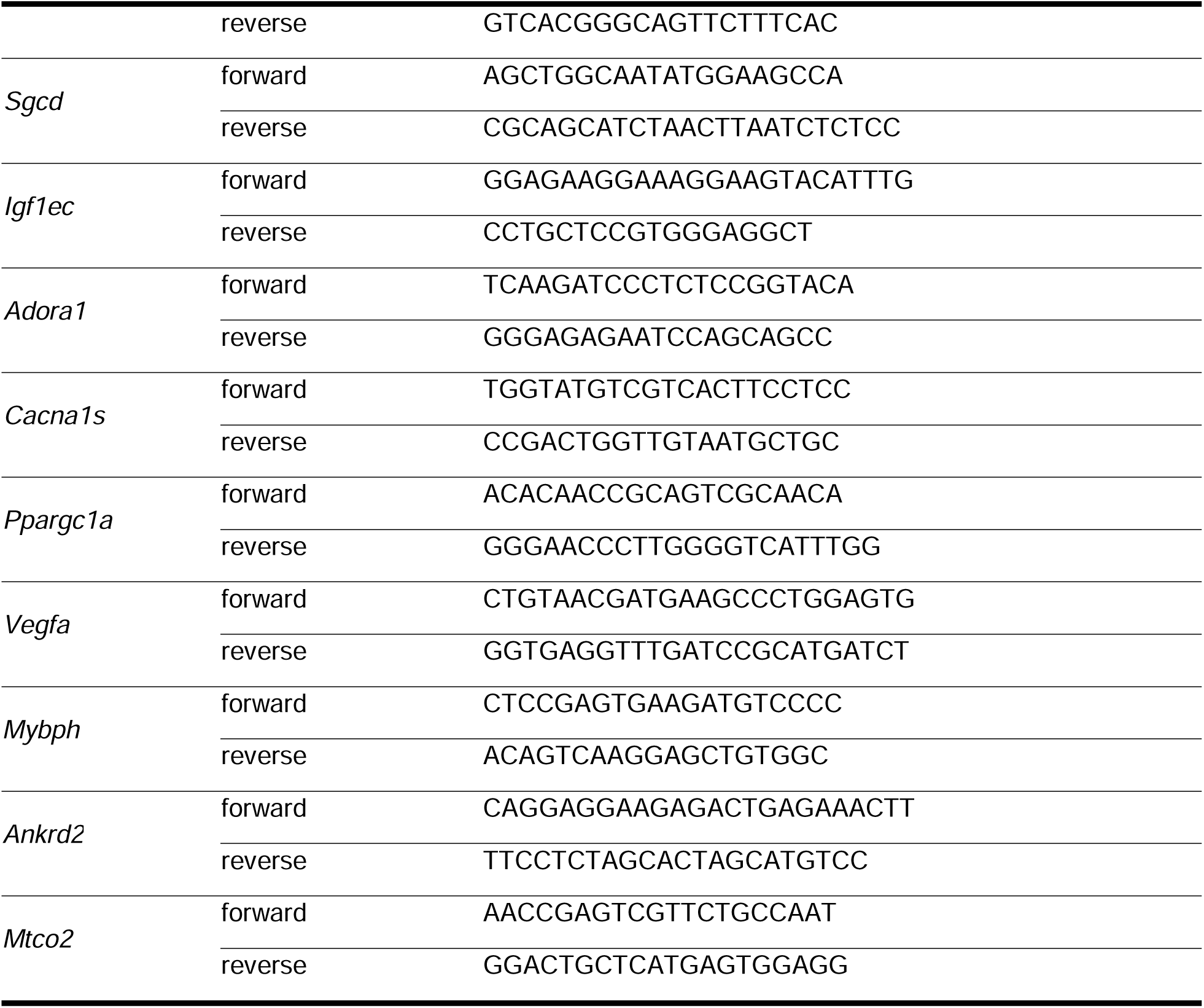
Primers sequences.

**Supplementary Data.**
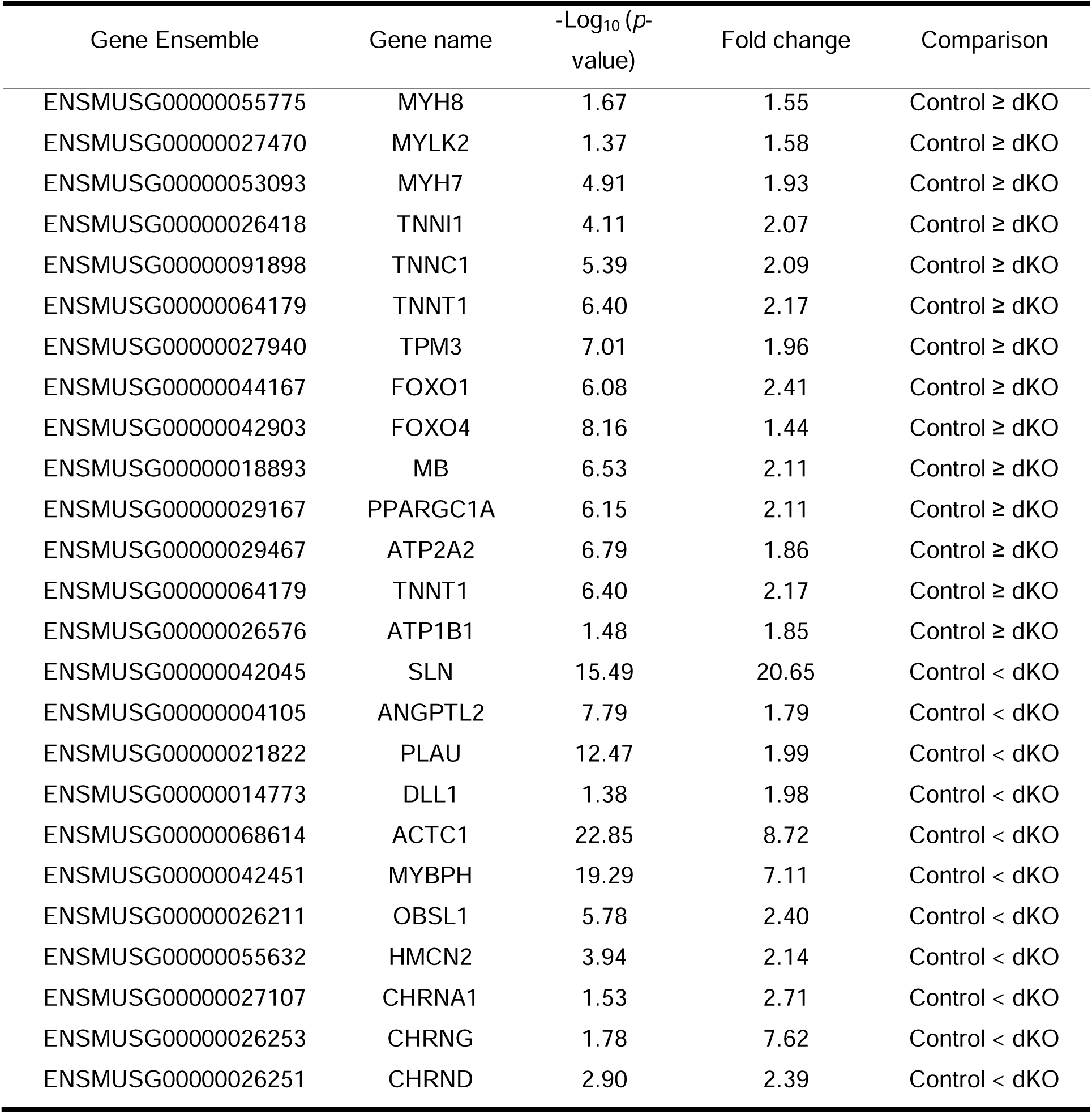
DEGs in gastrocnemius of dKO mice compared to control.

